# *FUS* ALS-causative mutations impact *FUS* autoregulation and the processing of RNA-binding proteins through intron retention

**DOI:** 10.1101/567735

**Authors:** Jack Humphrey, Nicol Birsa, Carmelo Milioto, David Robaldo, Andrea B Eberle, Rahel Kräuchi, Matthew Bentham, Agnieszka M. Ule, Seth Jarvis, Cristian Bodo, Maria Giovanna Garone, Anny Devoy, Alessandro Rosa, Irene Bozzoni, Elizabeth MC Fisher, Marc-David Ruepp, Oliver Mühlemann, Giampietro Schiavo, Adrian M Isaacs, Vincent Plagnol, Pietro Fratta

## Abstract

Mutations in the RNA-binding protein FUS cause amyotrophic lateral sclerosis (ALS), a devastating neurodegenerative disease in which the loss of motor neurons induces progressive weakness and death from respiratory failure, typically only 3-5 years after onset. FUS plays a role in numerous aspects of RNA metabolism, including mRNA splicing. However, the impact of ALS-causative mutations on splicing has not been fully characterised, as most disease models have been based on FUS overexpression, which in itself alters its RNA processing functions. To overcome this, we and others have recently created knock-in models, and have generated high depth RNA-sequencing data on FUS mutants in parallel to FUS knockout. We combined three independent datasets with a joint modelling approach, allowing us to compare the mutation-induced changes to genuine loss of function. We find that FUS ALS-mutations induce a widespread loss of function on expression and splicing, with a preferential effect on RNA binding proteins. Mutant FUS induces intron retention changes through RNA binding, and we identify an intron retention event in FUS itself that is associated with its autoregulation. Altered FUS regulation has been linked to disease, and intriguingly, we find FUS autoregulation to be altered not only by FUS mutations, but also in other genetic forms of ALS, including those caused by TDP-43, VCP and SOD1 mutations, supporting the concept that multiple ALS genes interact in a regulatory network.

## Introduction

Amyotrophic lateral sclerosis (ALS) is a relentlessly progressive neurodegenerative disorder characterised by loss of motor neurons, leading to muscle paralysis and death (Taylor, Brown, and Cleveland 2016). 5-10% of cases are inherited in an autosomal dominant fashion (Taylor, Brown, and Cleveland 2016). Numerous genes have been identified as disease-causative, and have been central to the understanding of pathogenesis. RNA-binding proteins (RBPs), most prominently TDP-43 and FUS, have been identified as a major category of causative genes in familial ALS (Sreedharan et al. 2008; Vance et al. 2009). In post-mortem brain tissue from mutation carriers, either TDP-43 and FUS are found to be depleted from the nuclei of cells, where they are normally predominantly localised. Instead both proteins are found to be accumulated in cytoplasmic inclusions, suggesting both a nuclear loss of function and a cytoplasmic gain of toxic function play a role in disease (Ling, Polymenidou, and Cleveland 2013). TDP-43 and FUS have multiple roles in RNA metabolism, including transcription, splicing, polyadenylation, miRNA processing and RNA transport (Polymenidou et al. 2011; Lagier-Tourenne et al. 2012; Rogelj et al. 2012; Ishigaki et al. 2012; Masuda et al. 2015; Ling, Polymenidou, and Cleveland 2013).

Although many studies have investigated the physiological functions of FUS and TDP-43 through knockout and overexpression experiments (Chiang et al. 2010; Iguchi et al. 2013; Wils et al. 2010; Shan et al. 2010; Wegorzewska et al. 2009; Barmada et al. 2010; Arnold et al. 2013; Hicks et al. 2000; Kino et al. 2015; Verbeeck et al. 2012; Mitchell et al. 2013; Shiihashi et al. 2016; Qiu et al. 2014; Sharma et al. 2016), the effect of disease-causing mutations on RNA splicing has been harder to investigate. Indeed, partly due to the fact that both proteins are very sensitive to dosage changes and very tightly regulated (Ayala et al. 2011; Zhou et al. 2013), mutation overexpression models are unfit to address these questions.

We and others have used mice carrying mutations in the endogenous *Tardbp* gene to show that TDP-43 mutations induce a splicing gain of function (Fratta et al. 2018; White et al. 2018). Here, we use our novel knock-in mouse model of FUS-ALS, FUS-Δ14 (Devoy et al. 2017), in combination with data from other physiological mouse and cellular models of FUS-ALS, to address the impact of ALS-causing FUS mutations on RNA metabolism, and splicing in particular. Although mutations have been observed throughout the FUS gene, the most aggressive FUS ALS-causing mutations cluster in the C-terminal region of the protein, where the nuclear localisation signal (NLS) resides (Figure 1A) (Shang and Huang 2016). These mutations affect the binding of the nuclear localisation signal by transportin and induce an increase in cytoplasmic localisation of the protein (Dormann et al. 2010; Shang and Huang 2016).

**Figure 1:**
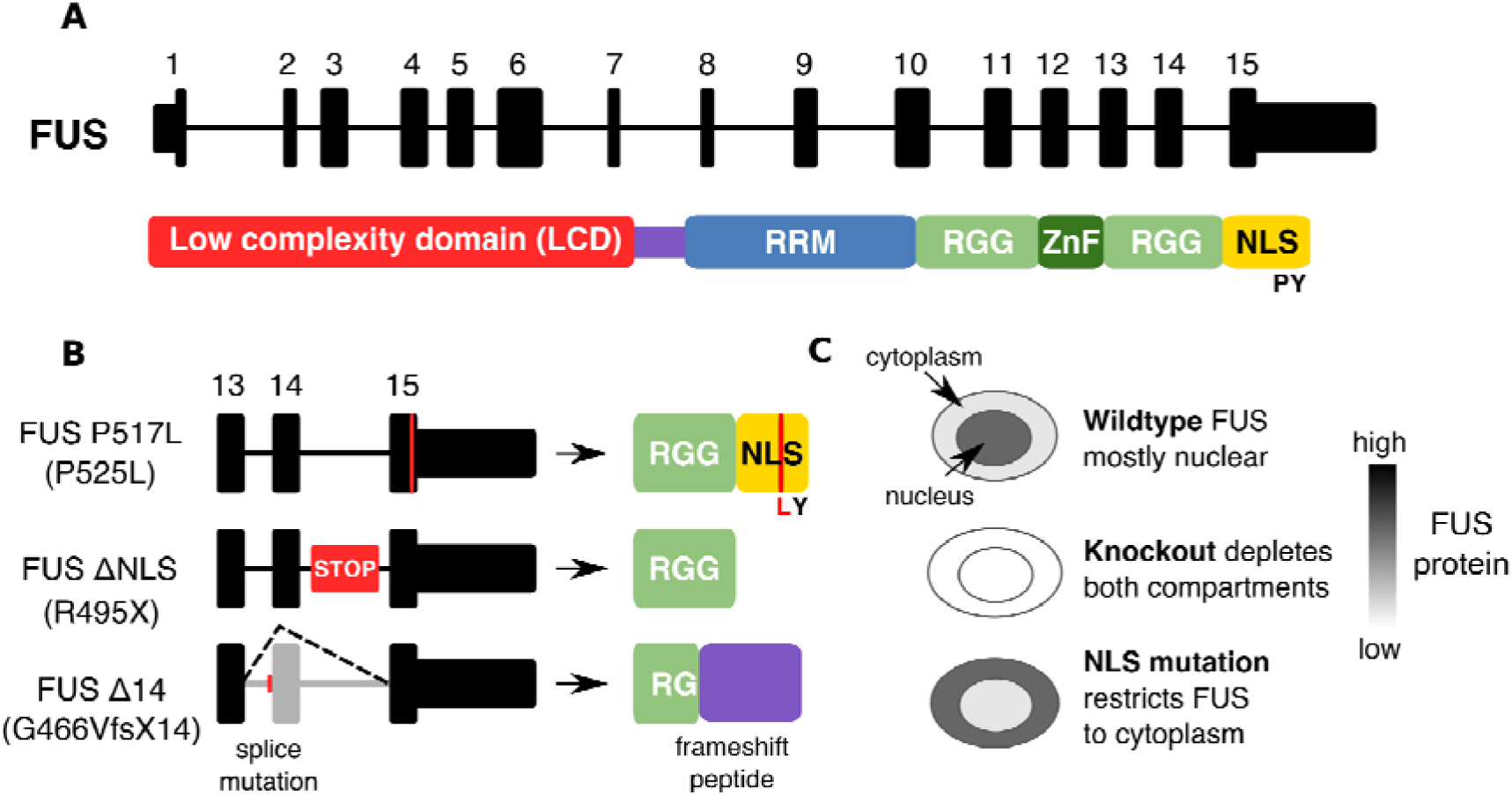
Illustration of the models and mutations used in this study. (**A**) The major transcript encoding the FUS protein in human and mouse is comprised of 15 exons. FUS protein is comprised of a low complexity domain (LCD), an RNA recognition motif (RRM) domain, two Arginine-Glycine-Glycine (RGG) domains, a zinc finger domain (Znf) and a nuclear localisation signal (NLS) (Shang and Huang 2016). (**B**) The three FUS NLS mutations used in this study. The Bozzoni group knocked in a point mutation to create the FUS P525L line, a missense mutation equivalent to the human ALS P517L mutation. The Dupuis group created a FUS ΔNLS line where the entire NLS is removed. We have used the FUS Δ14 mouse, where a frameshift mutation leads to the skipping of exon 14 and a frameshifting of the remaining NLS sequence. (**C**) In wildtype cells FUS protein is predominantly nuclear but can shuttle to the cytoplasm. When FUS is knocked out it will be reduced in both compartments but if the NLS is mutated or deleted then FUS will accumulate in the cytoplasm.

We find that FUS NLS mutations induce a splicing loss of function, particularly in intron retention events. These events are enriched in transcripts encoding RBPs, and FUS itself uses intron retention to autoregulate its own transcript. Finally, we show that this regulatory splicing event is altered in different ALS model systems. These findings shed light on primary changes caused by ALS mutations, on how RBPs act as a network to regulate each other, and on the mechanism for FUS autoregulation. This is of central importance for understanding the vicious cycle of cytoplasmic accumulation of RBPs driving further RBP expression and mislocalisation that ensues in disease (Fratta and Isaacs 2018) and is a target for therapeutic strategies.

## Results

### Joint modelling identifies high confidence FUS gene expression targets

To identify transcriptional changes induced by ALS-causing FUS mutations, we performed high depth RNA sequencing on spinal cords from FUS-Δ14 mice, a recently described knock-in mouse line carrying a frameshift mutation in the FUS C-terminus (Devoy et al. 2017). The mutation leads to a complete NLS loss (Figure 1A,B) and its replacement with a novel peptide. Analysis of homozygous mutations is important to identify the effects of mutant FUS and, as homozygous FUS-Δ14 mice are perinatally lethal, we used late-stage homozygous, heterozygous and wildtype (E17.5) embryonic spinal cords. In order to directly compare mutation-induced changes to genuine FUS loss of function, we performed similar experiments in parallel using FUS knockout (KO) and littermate control E17.5 spinal cords. We refer to these samples in the manuscript as the “Fratta samples”.

We took advantage of two publicly available mouse CNS datasets, where ALS-causative FUS mutations were inserted in the endogenous *Fus* gene, expressed homozygously, and where FUS KO was used in parallel, to identify changes relevant across FUS NLS mutations. The RNA-seq datasets (described in Supplementary Table 1A) were a) E18.5 brains from FUS-ΔNLS, a model of the R495X mutation that removes the entire NLS (Bosco et al. 2010), along with FUS-KO mice (Scekic-Zahirovic et al. 2016), together referred to as the “Dupuis samples”; and b) ES-derived motor neurons from FUS-P517L, corresponding to human P525L (Chiò et al. 2009) which mutates the critical proline residue of the NLS, along with a FUS-KO (Capauto et al. 2018), together referred to as the “Bozzoni samples” (Figure 1B). After performing differential expression analyses on each individual comparison, we combined the three datasets and performed two joint analyses for the KO and NLS mutation samples with their respective controls. Throughout the manuscript, we refer to these two joint models as KO and MUT respectively. This approach identifies differentially expressed and spliced genes that have a shared direction of effect between the three KO or MUT datasets.

At FDR < 0.05 the KO and MUT joint differential gene expression models contain 2,136 and 754 significantly differentially expressed genes respectively. When comparing the genes found by the two joint analyses to the six individual analyses there is only a moderate overlap (Supplementary Table 2), suggesting that a large number of genes called as significantly differentially expressed in a single comparison cannot be replicated in the others, despite being the same condition and all being generated from embryonic neuronal tissue.

### FUS NLS mutations have a loss-of-function effect on gene expression

We next looked for evidence of either a shared or divergent gene expression signal between the KO model and the MUT model. With a conservative threshold for overlap, where a gene must be significant at FDR < 0.05 in both models, we found an overlap of 425 shared genes between KO and MUT, with 329 genes being classified as mutation-specific and 1,711 as knockout specific. More permissive overlap criteria, where a gene overlaps if it reaches FDR < 0.05 in one model and an uncorrected P < 0.05 in the other, increased the overlap to 1,318 genes, reducing the specific genes to 186 in the MUT model, and 961 in KO (Figure 2A). Comparing the direction of changes found for the 1,318 overlapping genes between FUS KO and FUS MUT showed that only 7 genes are altered in opposing directions, confirming a loss of function effect of *Fus* mutations on gene expression (Figure 2B). Fitting a linear model between the fold changes of the two datasets showed that the effect of FUS MUT on gene expression is 76% that of FUS KO. (β = 0.76; P < 1e-16 F-test; R^2^ = 0.90). This suggests that while FUS KO and FUS MUT affect the same genes in the same directions, the magnitude of change is greater in FUS KO than FUS MUT. This relationship is not an artefact of the relaxed overlap criteria as fitting the model on just the 425 strictly overlapping genes had a similar outcome (β = 0.8; P < 1e-16). The relative weakness of NLS mutations compared to knockouts can be explained as NLS mutant FUS can still be detected in the nucleus, although at lower amounts (Devoy et al. 2017; Scekic-Zahirovic et al. 2017).

**Figure 2:**
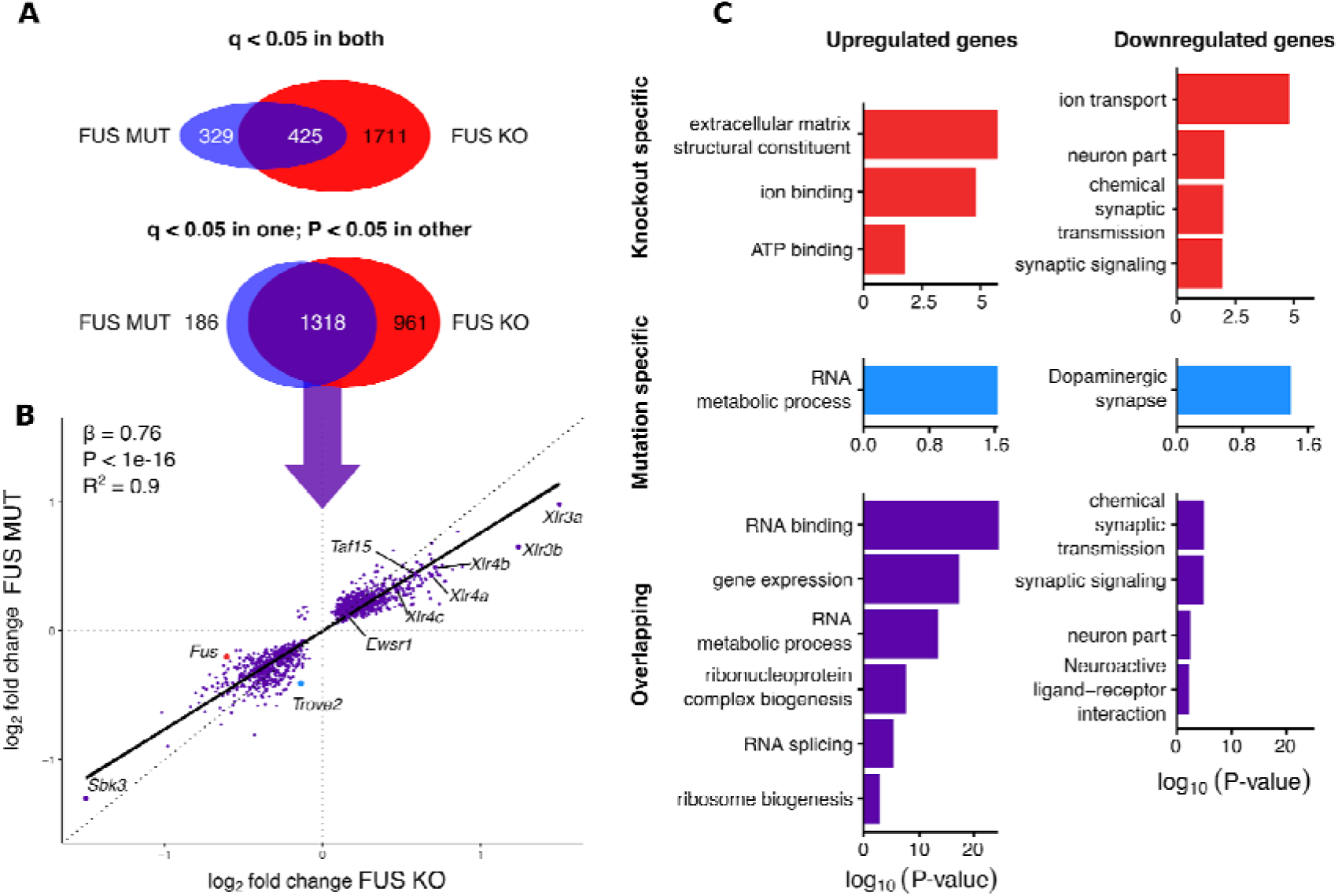
FUS mutations induce a loss of function on expression, with upregulation of RBPs and downregulation of neuronal genes. (**A**) Schematic of the strict and relaxed overlap thresholds between the two joint models of FUS KO and FUS MUT. (**B**) Plotting the log2 fold change for the MUT model against KO for the overlapping genes only, plus knockout-specific *Fus* and mutation-specific *Trove2*. (**C**) Gene Ontology terms enriched in the three categories of genes split by direction of change.

### FUS NLS mutations induce synaptic and RNA-binding gene expression changes

Amongst the most changed genes are the remaining members of the FET family of RNA-binding proteins, *Taf15* and *Ewsr1*, as well as *Trove2*, the gene encoding the 60 kDa SS-A/Ro ribonucleoprotein, which is downregulated in MUT only and unchanged in KO (Supplementary Figure 2). Interestingly, we found the X-linked mouse-specific lymphocyte receptor genes, *Xlr3a*, *Xlr3b*, *Xlr4a* and *Xlr4b* strongly upregulated in both conditions but more so in KO than MUT. These genes form a cluster of paralogous genes on the X chromosome and are paternally imprinted in mice (Raefski and O’Neill 2005). (Supplementary Figure 2).

Gene ontology (GO) analyses showed that genes commonly upregulated in both KO and MUT to be enriched in RNA binding, splicing and metabolism terms, and commonly downregulated genes in synaptic and neuronal terms (Figure 2C). KO-specific and MUT-specific genes were less clearly enriched in specific functions. (Figure 2C).

To investigate a relationship between FUS binding to particular genomic features and the direction of differential expression, we used a FUS iCLIP dataset from embryonic day 18 mouse brain (Rogelj et al. 2012). Intronic and 3’UTR binding was most common, as has been previously reported (Lagier-Tourenne et al. 2012; Rogelj et al. 2012; Ishigaki et al. 2012). In this analysis we compared differentially expressed genes to a non-differentially expressed set matched for gene length and expression and found enrichment of FUS binding specifically within downregulated genes. This effect was primarily driven by binding within introns (Supplementary Figure 2).

### FUS NLS mutations induce a splicing loss of function

We used the joint modelling approach to assess the impact of FUS MUT and KO on alternative splicing, including all possible alternative splicing isoforms, both novel and annotated, by combining the datasets together. The joint models increased power of detection of splicing changes for both MUT and KO, as respectively 93 and 890 events were found to be significantly altered, more than the sums of the individual analyses (Supplementary Table 3). There is also a very good concordance between each individual analysis and their joint model, with the exception of the Bozzoni dataset (only 7 out of 31). Comparison of splicing events between the heterozygous and homozygous FUS-Δ14 and FUS-KO samples found 34 overlapping FUS-Δ14 events and 115 overlapping FUS KO events (Supplementary Figure 3). Comparing fold changes showed heterozygotes to have a reduced effect size compared to the homozygotes in both FUS-Δ14 (beta = 0.57; P = 1e-11) and FUS KO (beta = 0.67; P < 1e-16), demonstrating a gene dosage effect on splicing.

Comparing the joint FUS KO and MUT splicing models, there are 405 overlapping events at a permissive significance threshold, with 501 KO specific splicing events and only 16 MUT-specific splicing events (Figure 3A). There are no overlapping splicing events that change in opposing directions, confirming FUS mutations have a loss of function effect on splicing. Furthermore, larger fold changes are present in the KO compared to the MUT joint models (β = 0.7, P < 1e-16; F-test; R^2^ = 0.89), supporting the reduced nuclear localisation of the FUS mutations as the main responsible factor for the loss of splicing function.

**Figure 3:**
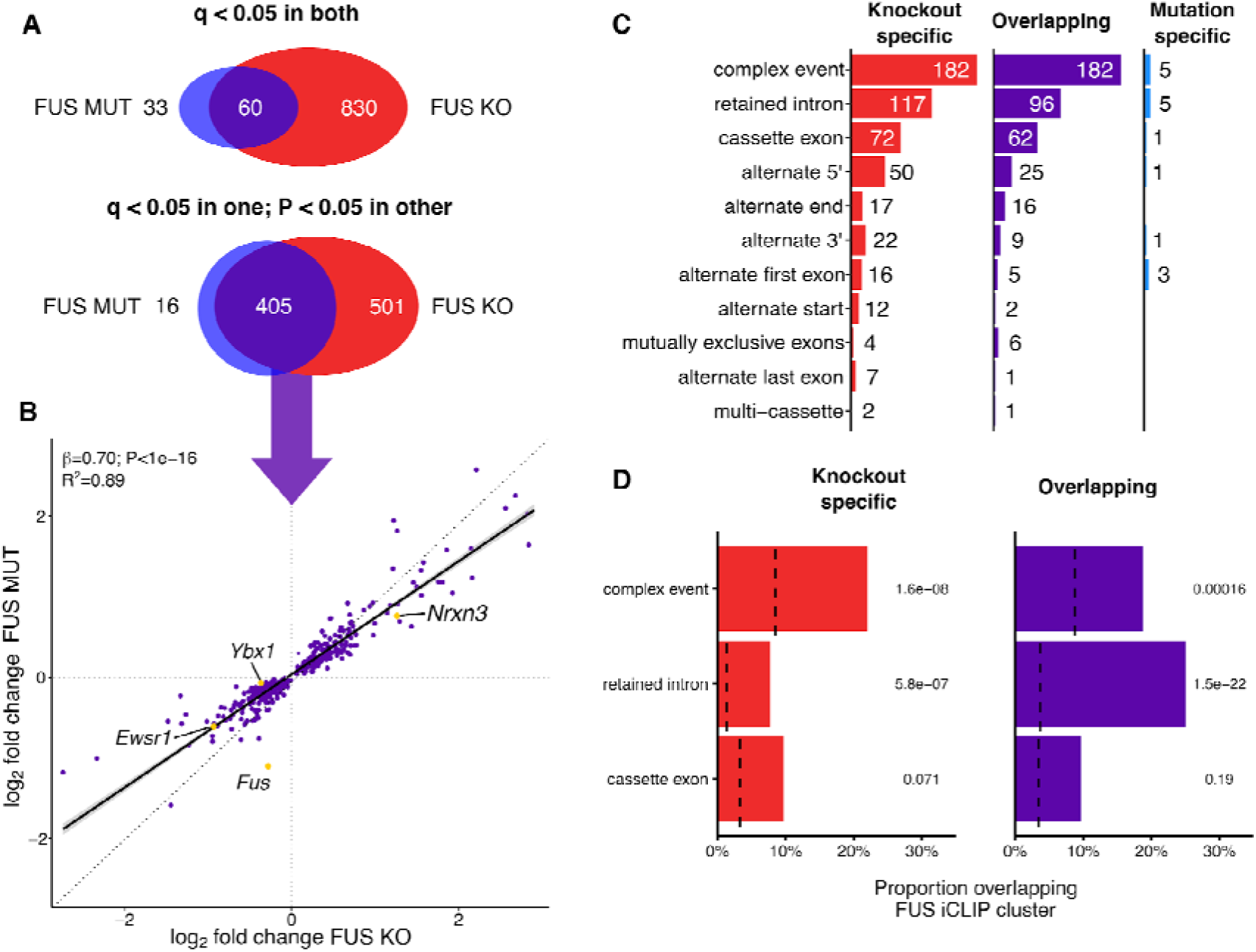
Mutant FUS induces a splicing loss of function. (**A**) Schematic of the strict and relaxed overlap thresholds between the two joint splicing models of FUS KO and FUS MUT. (**B**) Plotting the log2 fold change for the MUT model against KO for the overlapping splicing events. (**C**) Counts of each category of splicing events found in the three sets. (**D**) The proportion of each type of splicing variant in each category that overlap a FUS iCLIP cluster. Background sets of non-regulated splicing events matched for length and wildtype expression are represented by dotted lines. P-values from proportion test, corrected for multiple testing with the Bonferroni method.

### Intron retention is the most common splicing change induced by FUS mutations

Separating splicing events by type showed a similar distribution between KO-specific and MUT/KO overlapping events as both are dominated by retained introns and complex events (Figure 3C). The latter are difficult to interpret as multiple types of alternative splicing co-occur within the same locus. This can be seen in *Ybx1*, where a retained intron is accompanied by an alternate cassette exon and the splicing of both are altered in both FUS KO and MUT (Supplementary Figure 3). Cassette exons and alternate 5′ and 3′ splice sites were found in all three sets of genes, with alternate 5′ sites appearing at twice the rate of alternate 3′ splice sites. FUS has been shown to interact with the U1 snRNP, which may explain this over-representation (Yu et al. 2015; Yu and Reed 2015).

Methods that observe RNA-protein interactions have proposed direct regulation of splicing by FUS through binding to introns (Ishigaki et al. 2012; Lagier-Tourenne et al. 2012; Rogelj et al. 2012; Masuda et al. 2015). We used published FUS iCLIP clusters (Rogelj et al. 2012) to show that retained introns are strongly enriched for FUS binding sites, with the strongest enrichment present for the overlapping MUT/KO retained introns (P = 4.4e-22; Chi-squared; Figure 3D). No enrichment was seen in cassette exons, suggesting that these events may not be the direct result of altered FUS binding.

### FUS-regulated retained introns are enriched in RBPs and are highly conserved

Gene ontology analysis for each category of events showed a clear enrichment in RNA-binding and neuronal GO terms in the overlapping splicing events. Specifically, genes with retained introns were often related to RNA binding (Figure 4A). The RNA-binding transcripts include the U1 splicing factor *Snrnp70,* the FET protein family members *Ewsr1* and *Taf15,* and *Fus* itself. Conversely, neuronal terms were only enriched in cassette exons.

**Figure 4:**
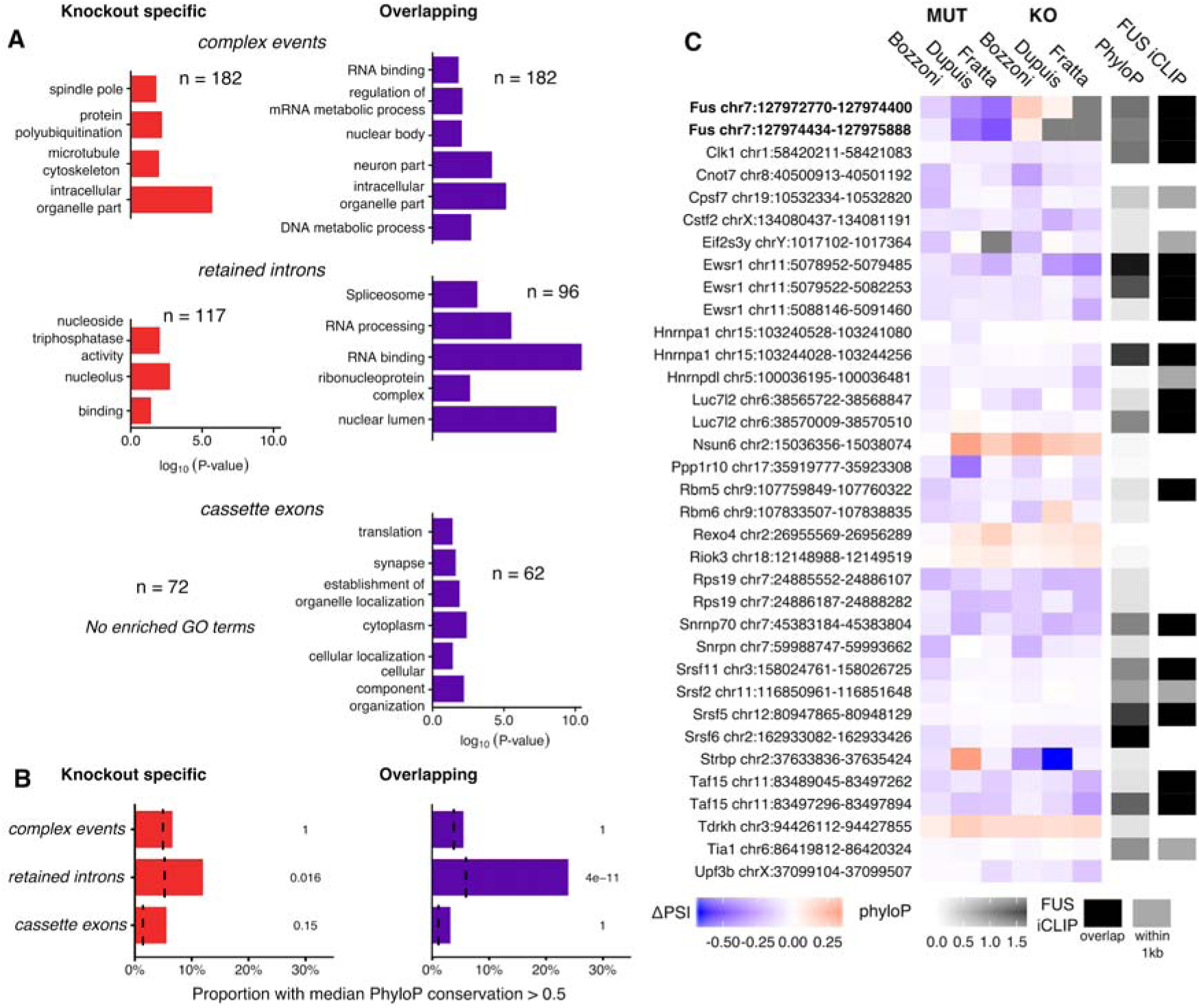
FUS modulates the inclusion of a set of highly conserved RNA-binding protein introns. (**A**) Significantly enriched Gene Ontology terms found in genes split by category and splicing variant type. (**B**) The proportion of each type of splicing event that has a median phyloP conservation score greater than 0.5. Background sets as before. P-values from proportion test, corrected for multiple testing with the Bonferroni method. (**C**) All intron retention events found in the Overlapping set found to have an RNA-binding GO term, along with the two FUS introns which are mutation-specific. PSI Δ values were calculated for each individual splicing analysis and presented from negative (blue) to positive (red). Events not identifiable in a dataset are coloured grey. Median phyloP conservation across each intron coded from 0 (non-conserved; white) to 1.5 (highly conserved; black). Additionally, each intron is noted for the presence of FUS iCLIP cluster overlapping (black) or within 1kb of either end of the intron (dark grey).

RNA-binding proteins often contain intronic sequences that are very highly conserved (Lareau et al. 2007) and have been proposed to be important for their post-transcriptional regulation (Ni et al. 2007). To test whether the MUT FUS-targeted splicing events show high sequence conservation, we calculated the median phyloP score using the 60-way comparison between mouse and other species for each encompassing intron (Pollard et al. 2010). Sets of events were then tested on the proportion of the set with a median phyloP score >0.5, where a score of 0 is neutral and >1 is highly conserved. Only retained introns were enriched in sequence conservation, and at a greater extent for MUT/KO overlapping (P = 1.2e-10) than KO-specific events (P = 0.048; Figure 4B). 35 retained intron events are found in genes with RNA-binding GO terms. Calculating the direction of change shows these introns to be predominantly decreasing in retention upon FUS KO or NLS mutation. In addition, 20/35 have a FUS iCLIP cluster within 1kb of the intron (Figure 4C).

Taken together, these results show that nuclear loss of FUS through either knockout or NLS mutation leads to a set of splicing changes enriched in conserved intron retention events affecting predominantly RNA-binding proteins. Conversely, cassette exons are not bound by FUS beyond the null expectation and originate from nonconserved introns.

### FUS autoregulates through highly conserved retained introns

The joint splicing analyses found two retained introns (introns 6 and 7) in the *Fus* transcript to be less retained in FUS NLS mutants. Both introns are highly conserved and contain large numbers of FUS iCLIP peaks (Figure 5A; Supplementary Figure 4). Numerous RBPs regulate their expression by binding their own transcript (Dredge et al. 2005; Rossbach et al. 2009; Wollerton et al. 2004; Ayala et al. 2011). Through this autoregulation, when protein levels are high, increased binding of the pre-mRNA shifts alternate splicing towards the production of an untranslated isoform, either by including a premature stop codon, which is sensitive to nonsense-mediated decay (NMD), or by detaining the transcript in the nucleus to avoid translation (Rosenfeld, Elowitz, and Alon 2002; Jangi et al. 2014; McGlincy and Smith 2008; Boutz, Bhutkar, and Sharp 2015). The FUS intron 6/7 region is the putative locus of FUS autoregulation through the skipping of exon 7, which causes a frameshift, producing an NMD-sensitive transcript (Zhou et al. 2013). However, when examining RNA-seq junction coverage of the FUS gene in all our mouse datasets, we failed to observe skipping of exon 7 in any sample. Instead, the retention of both introns 6 and 7 decreased in the presence of FUS mutations in all three datasets (Figure 5B), despite the baseline level of intron retention in wildtype samples being highly variable between datasets. Heterozygous FUS-Δ14 mice also showed a significant reduction in intron retention, albeit less than the homozygotes, demonstrating a dose-dependent response. We validated our findings using RT-PCR and confirmed that intron retention decreases in a mutation dose-dependent manner (Figure 5C; intron 6 P = 5.7e-4; intron 7 P = 8.7e-4; ANOVA). We failed to detect a band corresponding to the skipping of exon 7 in any sample.

**Figure 5:**
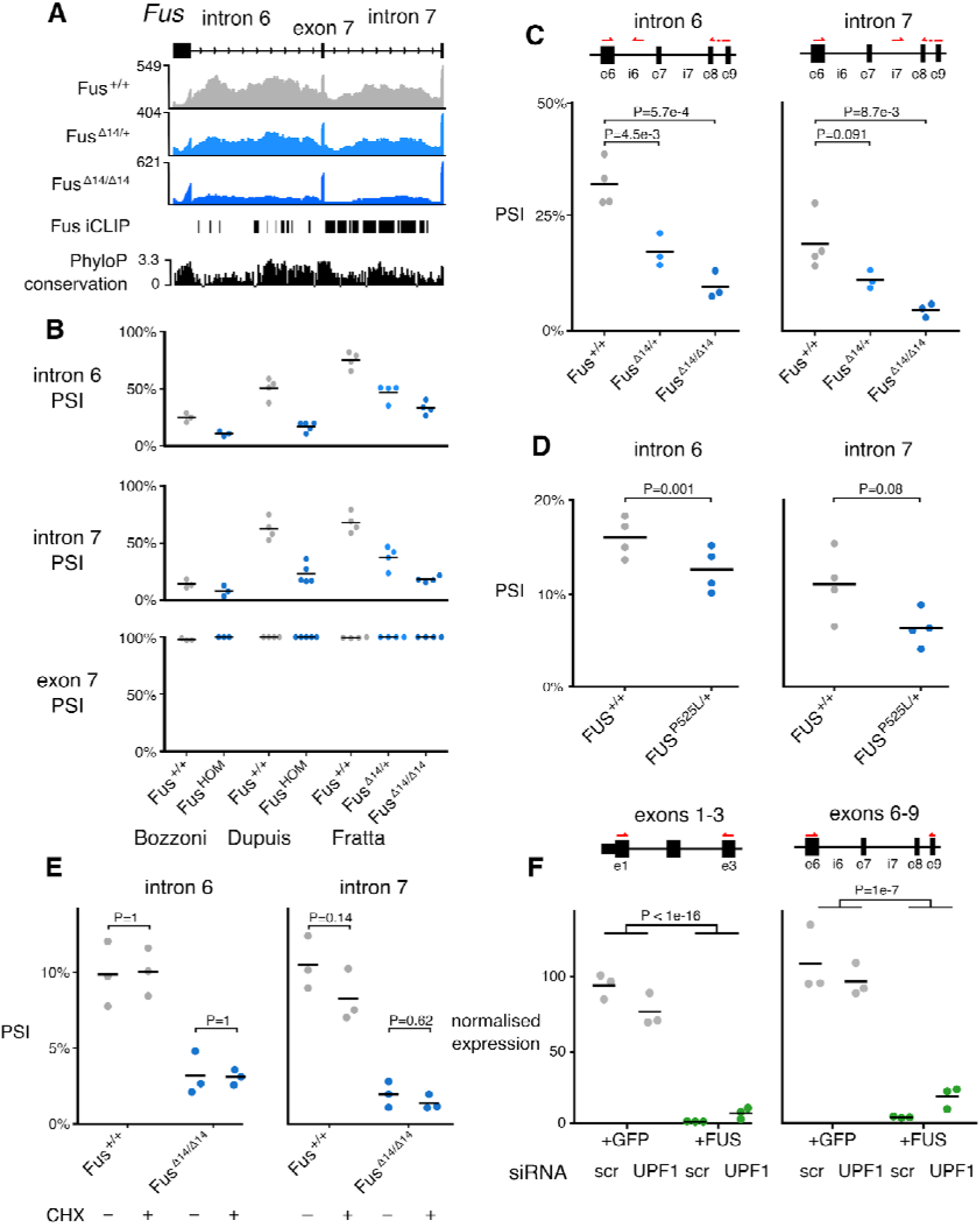
FUS autoregulation is dependent on intron retention. (**A**) FUS introns 6 and 7 are highly conserved and have multiple FUS iCLIP binding peaks. Retention of introns 6 and 7 decreases with increasing dose of FUS Δ14. RNA-seq coverage for wildtype, FUS Δ14 heterozygous and FUS Δ14 homozygous samples are accompanied by FUS iCLIP (Rogelj et al., 2012) and phyloP conservation (60 way) tracks. (**B**) Percentage spliced in (PSI) values of intron 6, intron 7 and exon 7 in the three datasets, including the FUS Δ14 heterozygotes. FUS^HOM^ refers to homozygous FUS P517L in the Bozzoni dataset, and homozygous FUS ΔNLS in Dupuis. (**C**) RT-PCR validation of the reduction in intron 6 and 7 inclusion with increasing dose of FUS Δ14 mutation. Left pane: FUS intron 6; ANOVA genotype P=5.1e-4. Right panel: FUS intron 7; ANOVA genotype P=8.5e-3. Pairwise t-tests reported on plot, corrected by Holm method. (**D**) RT-PCR validation of reduced retention of FUS introns 6 and 7 in fibroblasts from human patient with a FUS P525L mutation (n=1) compared to a healthy control (n=1). RT-PCR repeated in triplicate for each sample. (**E**) Translation blocked with cycloheximide (CHX) to observe whether the intron retention transcript is sensitive to nonsense-mediated decay. Left panel: FUS intron 6 retention is not altered with CHX treatment. ANOVA treatment P =0.96; genotype P =5.7e-5; interaction P = 0.86. Right panel: FUS intron 7 retention is unchanged by CHX treatment. ANOVA treatment P=0.10; genotype P = 7.9e-6; interaction P = 0.1. Pairwise t-tests reported on plot, corrected by Holm method. (**E)** RT-PCR on FUS introns 6 and 7 repeated on nuclear and cytoplasmic fractions. Left panel: intron 6 retention is reduced in the cytoplasm of FUS mutants. ANOVA fraction P = 0.01; genotype P = 0.001; interaction P = 0.02. Right panel: intron 7 retention is reduced in the cytoplasm in both wildtype and FUS mutant cells. ANOVA fraction P = 0.006; genotype P = 2.4e-4; interaction P = 0.77. Pairwise t-tests reported on plot, corrected by multiple testing by Holm method. **(F)** Reduced endogenous FUS RNA levels HeLa cells expressing codon-optimised FUS compared to HeLa cells expressing GFP as a control, as measured by qPCR (n=3). This reduction was mostly unaffected by the siRNA depletion of UPF1.

We then tested whether the same phenomenon of reduced intron retention occurs in human cells, and used primary fibroblasts from a patient carrying the ALS-causative heterozygous FUS mutation G496Gfs that induces a strong cytoplasmic FUS mislocalisation (Devoy et al. 2017). RT-PCR showed a decrease in both intron 6 and 7 retention relative to a FUS wildtype human sample (Figure 5D; intron 6 P = 0.001; intron 7 P = 0.08; ANOVA). These changes were smaller than those observed in homozygous mice, and more similar to the heterozygous mice.

In order to further validate FUS autoregulation, we investigated the effect of FUS overexpression. FUS overexpression has previously been shown to downregulate endogenous *Fus* transcripts (Mitchell et al. 2013; Zhou et al. 2013; Ling et al. 2019). However, the presence of an overexpression plasmid makes it difficult to evaluate the expression of endogenous *FUS*. To circumvent this, we generated FlpIn HeLa cell lines that constitutively overexpress *FUS* cDNA using alternative codons, allowing us to design qPCR primers that selectively amplify endogenous *FUS*, but not the overexpressed version. We investigated the effects of FUS overexpression on endogenous *FUS* expression using qPCR. As expected, FUS overexpressing cells showed a strong downregulation of endogenous *FUS* mRNA compared to cells expressing a GFP construct (Exons 1-3 P < 1e-16; Exons 6-9 P = 1e-7; ANOVA; Figure 5F).

### Fus intron 6/7 retention determines transcript nuclear detention

Retaining two introns would be expected to cause NMD through the presence of premature stop codons, which are abundant in both intron 6 and 7 in all frames. We performed the mouse *Fus* intron 6/7 RT-PCR following incubation with cycloheximide (CHX) to block translation, thereby inhibiting NMD. We observed no change in intron retention in either wildtype or FUS-Δ14 homozygous cells (Figure 5E), despite observing a robust inhibition of NMD when looking at a known event in Srsf7 (Supplementary Figure 7). Similar results were obtained in human cells when UPF1, an essential NMD factor, was knocked down using siRNA (Figure 5F), showing only a modest effect on FUS mRNA levels. Taken together, these experiments show that *FUS* intron retention is NMD-insensitive.

Retained intron transcripts have been shown to accumulate in the nucleus (Boutz, Bhutkar, and Sharp 2015), thereby avoiding NMD, which occurs in the cytosol following RNA export. FUS introns 6 and 7 have been observed to be retained in the nucleus in human cells in a study using the APEX-seq method to label RNA in different cellular compartments (Fazal et al. 2018). We therefore suggest that FUS regulates its own expression through the retention of introns 6 and 7. This transcript is most likely detained in the nucleus, reducing the amount of cytoplasmic FUS available for translation.

### FUS intron retention is co-regulated by TDP-43 and altered in other ALS models

FUS shares RNA targets with another ALS-associated RNA-binding protein, TDP-43 (Lagier-Tourenne et al. 2012). Despite FUS iCLIP peaks being present throughout the *Tardbp* gene, neither FUS knockout or NLS mutation altered *Tardbp* expression or splicing.

However, using TDP-43 CLIP data from human and mouse collected by the POSTAR database (Hu et al. 2017), we identified a conserved TDP-43 binding site within *FUS* intron 7, approximately 400 nucleotides downstream of the 5’ splice site in a GU-rich region conserved between mouse and humans (Figure 6A,B). GU dinucleotides are the known binding motif of TDP-43 (Lukavsky et al. 2013). To test whether TDP-43 is also involved in regulating the retention of FUS introns 6 and 7, we re-analysed RNA-seq data where TDP-43 was knocked down in adult mice with an antisense oligonucleotide (Polymenidou et al. 2011) as well as adult mice homozygous for a C-terminal TDP-43 mutation (M323K) which we previously showed to cause a gain of splicing function (Fratta et al. 2018). TDP-43 knockdown caused a reduction in FUS intron retention, similar to FUS NLS mutations (Figure 6C). Conversely, TDP-43 M323K mice had an increase in FUS intron retention relative to wildtype. We observed no changes in FUS intron retention in embryonic mice homozygous for a splicing-null mutation (F210I) (Fratta et al. 2018), suggesting a developmental component to TDP-43 cross-regulation of FUS. FUS introns 6 and 7 are also bound by fellow FET family members TAF15 and EWSR1 in human and mouse (Supplementary Figure 4).

**Figure 6:**
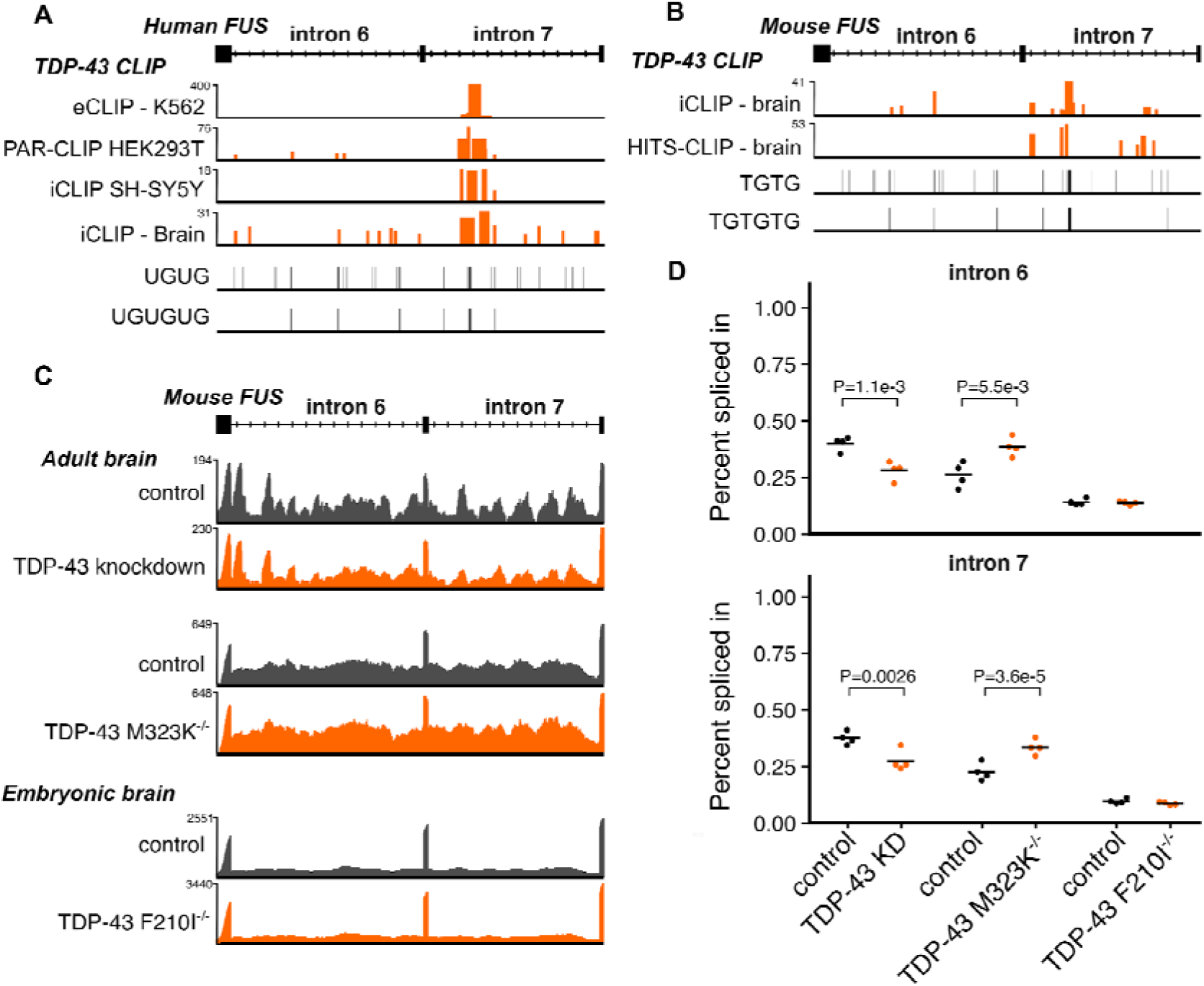
TDP-43 co-regulates FUS intron retention. (**A**) Mouse TDP-43 cross-linking and immunoprecipitation (CLIP) data overlap a TG-rich section of Fus intron 7. (**B**) Human TDP-43 CLIP data overlap within FUS intron 7 in a section rich with TG sequence. (**C**) RNA-seq traces of representative samples demonstrate decreased *Fus* intron retention in TDP-43 knockdown and increased retention in TDP-43 M323K mutation, both in adult mouse brain. No effect is seen with the RNA-binding mutant F210I in embryonic mouse brain. Y axis of each trace refers to the maximum read depth. (**D**) Percentage spliced in quantification from each TDP-43 dataset of intron 6 and 7 retention. P-values are presented from splicing analysis on each dataset.

**Figure 7:**
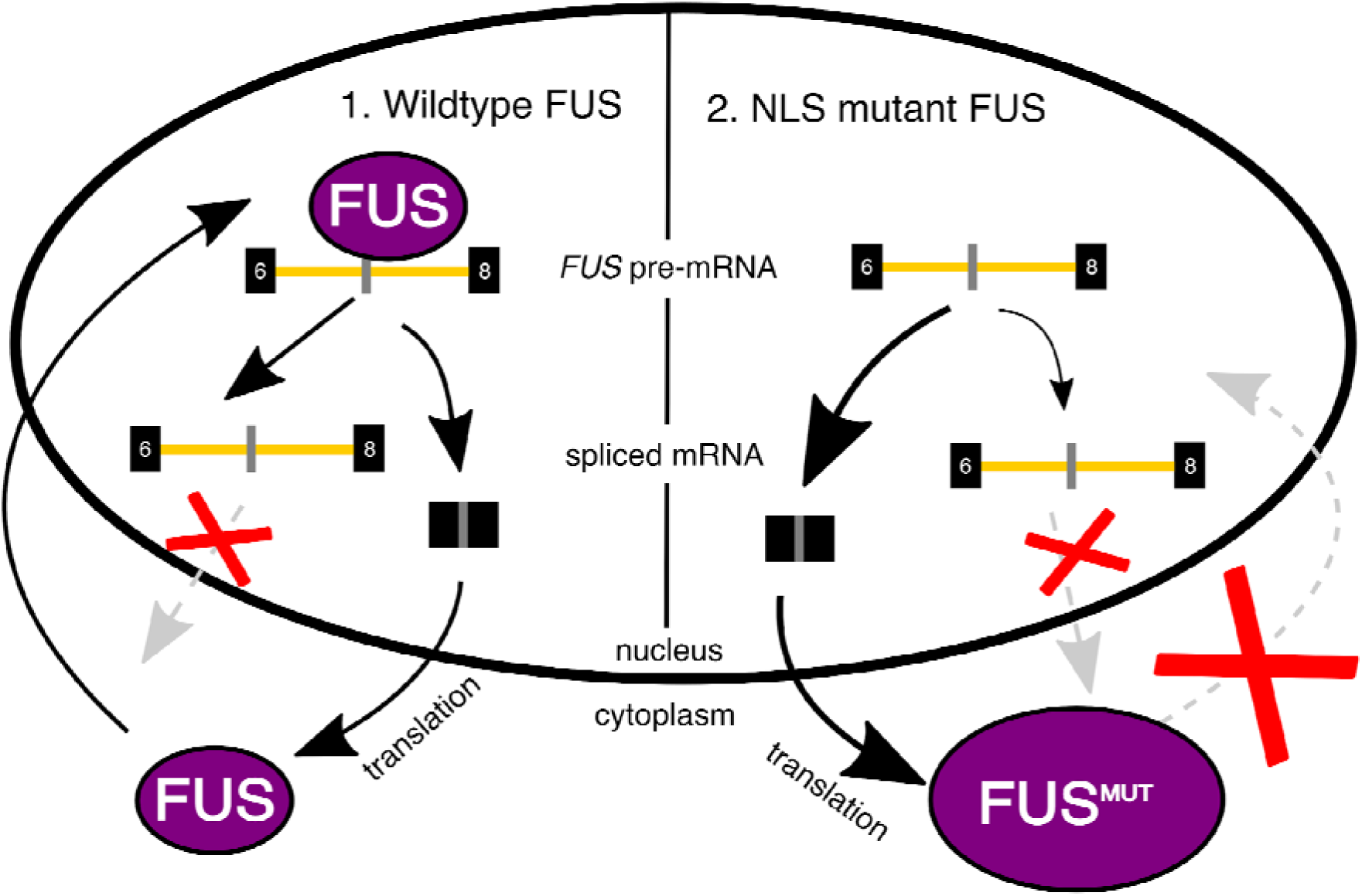
Cartoon summary. In wildtype cells FUS protein shuttles between the nucleus and cytoplasm. FUS binding within *FUS* introns 6 and 7 promotes their retention. The intron retention transcript is restricted to the nucleus, reducing the amount of cytoplasmic *FUS* mRNA available for translation. In conditions of low FUS protein, intron retention will be reduced and cytoplasmic *FUS* transcript will be increased. In contrast, in cells with FUS NLS mutations, mutant FUS is not transported to the nucleus as effectively. This reduces the ability of FUS protein to regulate FUS mRNA production through intron retention. This could lead to a vicious cycle of ever-increasing FUS protein in the cytoplasm, which may have toxic effects.

A previous study differentiated human motor neurons from induced pluripotent stem cells (iPSC) with and without mutations in VCP, an ALS gene that induces TDP-43 pathology (Hall et al. 2017; Luisier et al. 2018). The authors observed a set of retained intron events that change earlier in differentiation in the VCP mutant cells compared to the wildtype. These changes occurred primarily in RNA-binding proteins, suggesting a perturbation in splicing factor networks during development. To compare the FUS-regulated murine splicing events with those found in VCP mutant human cells, we overlapped the 143 human genes found to have intron retention events with the set of 219 mouse genes with either intron retention or complex events in both FUS MUT and FUS KO and found an overlap of 12 genes (P = 1e-5, Fisher’s exact test; Supplementary Table 4). Remarkably this included FUS itself. Re-analysis of published RNA-seq data from Luisier et al. demonstrated that the retention of FUS introns 6 and 7 is increased specifically in the transition between iPSC and neural precursor cell (NPC) stage in the VCP mutants (Supplementary Figure 8A).

Luisier and colleagues also found intron retention changes in another ALS model, SOD1 A4V mutations in iPSC-derived motor neurons (Kiskinis et al. 2014). Re-analysis of the RNA sequencing data showed a robust increase in FUS intron retention in the presence of SOD1 mutations (Supplementary Figure 8B). These findings suggest that FUS regulation is altered in the presence of ALS-causative mutations in other genes.

## Discussion

FUS is central player in ALS biology and its role in RNA metabolism has been intensively studied, but how disease-causing mutations impact upon RNA processing is still to be determined. One limitation has been that FUS, like many other RBPs, is extremely sensitive to gene dosage. Therefore, commonly used models, where mutant FUS is overexpressed, cannot disentangle the effects of overexpression from those of the mutation. We and others have generated FUS-ALS models where mutations were inserted into the endogenous gene to observe the impact on gene expression and splicing. In order to specifically investigate splicing changes, we generated high depth and long-read sequencing data from spinal cords of our mutant FUS mice, alongside FUS knockout samples, to compare mutant-induced changes to a pure loss of function. In order to identify with high confidence changes relevant to multiple FUS-ALS models, we performed a joint analysis of our data along with other publicly available datasets where endogenous *Fus* mutations had been studied in parallel to samples from knockout tissue. The sequencing conditions differed between datasets (Supplementary Table 1B), with some more suited for expression analysis rather than for splicing analysis. Nonetheless, the joint analysis model proved to be extremely powerful and allowed us to identify more splicing changes than using any single dataset independently. Furthermore, this approach limits artifactual findings from single datasets and allowed us to define a comprehensive high-confidence list of both expression and splicing targets induced by ALS-FUS mutations and FUS loss.

The comparisons between the joint analysis of FUS mutations and FUS knockout show that FUS mutations have a loss of function effect both on expression and splicing. This is a key difference from mutations in TDP-43, the other major RBP implicated in ALS, where specific mutations lead to gain of splicing function (Fratta et al. 2018; White et al. 2018). The effect appears weaker in NLS mutations than in complete knockout. This is compatible with the fact that mutant FUS is still found at low levels in nuclei of all the analysed mutants.

The high depth and long reads in our sequencing data allowed us to conduct an unbiased analysis of splicing, which highlighted intron retention and complex splicing events to be the most frequently altered class of changes, whilst cassette exon events, which had been previously described by using a targeted approach (Scekic-Zahirovic et al. 2016), are less abundant. Interestingly, when we assessed the link of FUS binding to different splicing events, FUS was linked directly to intron retention, but not to cassette exon splicing, suggesting the latter category could be due to downstream or indirect effects. Indeed, genes changed both at the splicing and expression level by *Fus* mutations are enriched in “RNA metabolism” GO terms, supporting a secondary effect on splicing of other RBPs.

Intriguingly, the FUS transcript features amongst the RBPs with intron retention changes induced by FUS mutations, with the mutation inducing a reduction in retention of introns 6 and 7. This region had been previously suggested to be involved in FUS autoregulation through alternative splicing of exon 7 (Zhou et al. 2013), but we were unable to observe this at significant levels in any of our datasets. The two introns are highly conserved across species, and FUS iCLIP data showed widespread FUS binding across both introns, supporting their retention as a putative autoregulatory mechanism. Although from looking at the translated sequence the FUS retained introns would be predicted to undergo NMD, we instead found the retained intron transcripts to be NMD-insensitive. Instead we find they could be detained in the nucleus (Boutz, Bhutkar, and Sharp 2015), a phenomenon also observed in a recent study on RNA localisation within different cellular compartments (Fazal et al. 2018) (**Figure 8**). The discrepancy between our results and those of Zhou and colleagues could be explained by FUS autoregulation having two independent mechanisms, using either detained introns and/or NMD-sensitive exon skipping, depending on the cell type and developmental stage. This would be akin to TDP-43, where both the nuclear retention of a long 3’UTR transcript has been observed, in addition to the production of an NMD-sensitive transcript through 3’UTR splicing (Ayala et al. 2011; Koyama et al. 2016). Further work could elucidate which autoregulatory mechanism predominates in affected motor neurons of FUS-ALS patients.

The observation that FUS mutations induce changes in other RBPs is compatible with the growing evidence of RBPs functioning as a sophisticated regulatory network (Jangi et al. 2014; Mohagheghi et al. 2016; Huelga et al. 2012; Gueroussov et al. 2017). This raises the question as to what other proteins contribute to the regulation of FUS levels, and whether other ALS-linked proteins could play a role. FUS binds and regulates the levels and splicing of EWSR1 and TAF15, both associated with ALS through rare familial mutations (Ticozzi et al. 2011; Couthouis et al. 2012) and both EWSR1 and TAF15 bind to *Fus* introns 6/7 (Supplementary Figure 4). Although FUS depletion leads to an upregulation of TAF15, reducing TAF15 has no effect on FUS expression (Kapeli et al. 2016). Furthermore, we found TDP-43 to bind reproducibly to intron 7 in both human and mouse, and that in adult mice TDP-43 knockdown induces a significant decrease in retention of both introns, whilst the opposite was found in the presence of gain-of-function TDP-43 mutations. No changes were found in an embryonic dataset from mice where TDP-43 has decreased RNA binding capacity. This observation could be due to the fact that TDP-43 is expressed at extremely high levels at the embryonic stage, and potentially relies on separate regulatory systems in development (Sephton et al. 2010; Cragnaz et al. 2015). Lastly, *FUS* intron retention is also altered in the presence of both VCP and SOD1 mutations, two ALS-related genes that are not RBPs and have recently been linked to intron retention (Luisier et al. 2018).

Of note, ALS forms have been subdivided by the pathology findings and FUS-ALS, TDP-43 ALS (which includes VCP-ALS) and SOD1 ALS, although clinically similar, show mutually exclusive proteins to be accumulated in post-mortem brain, raising the possibility they could act through independent mechanisms. Our findings showing FUS regulation to be altered by TDP-43, VCP and SOD1 mutations suggest that these proteins act upon a common network in disease.

In conclusion, we have found that FUS mutations induce a loss of splicing function, particularly affecting intron retention events in other RBPs. We show that an intron retention event in the *FUS* transcript is a putative mechanism for its autoregulation and is modified not only by mutations in FUS, but also by ALS-causative mutations in TDP-43, VCP and SOD1. These findings inform the understanding of both FUS-ALS and the wider biology of ALS.

## Supporting information

joint model expression results

GO terms for differentially expressed genes

joint model splicing results

GO terms for differentially spliced genes

---

**Supplementary Table 1A:**
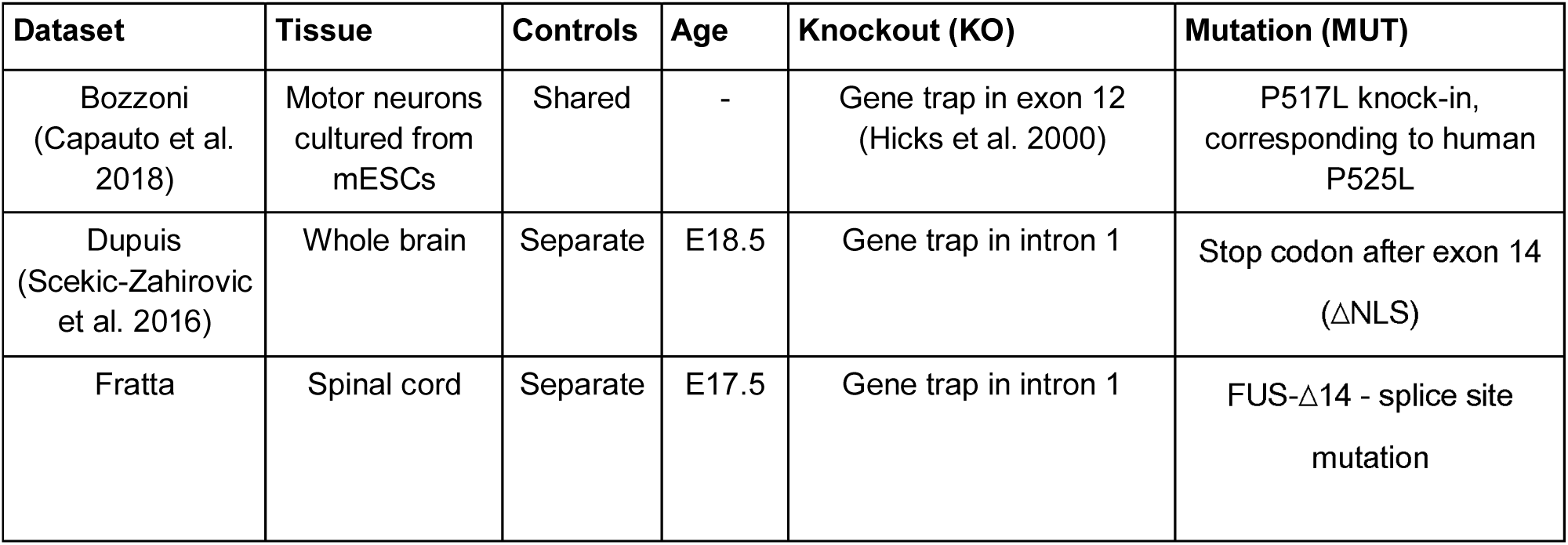
The three FUS mouse datasets used in this study. mESC - mouse embryonic stem cell

| Dataset | Tissue | Controls | Age | Knockout (KO) | Mutation (MUT) |
| --- | --- | --- | --- | --- | --- |
| Bozzoni<br>(Caputo et al. 2018) | Motor neurons cultured from mESCs | Shared | - | Gene trap in exon 12 (Hicks et al. 2000) | P517L knock-in, corresponding to human P525L |
| Dupuis<br>(Scekic-Zahirovic et al. 2016) | Whole brain | Separate | E18.5 | Gene trap in intron 1 | Stop codon after exon 14 (ΔNLS) |
| Fratta | Spinal cord | Separate | E17.5 | Gene trap in intron 1 | FUS-Δ14 - splice site mutation |

**Supplementary Table 1B:**
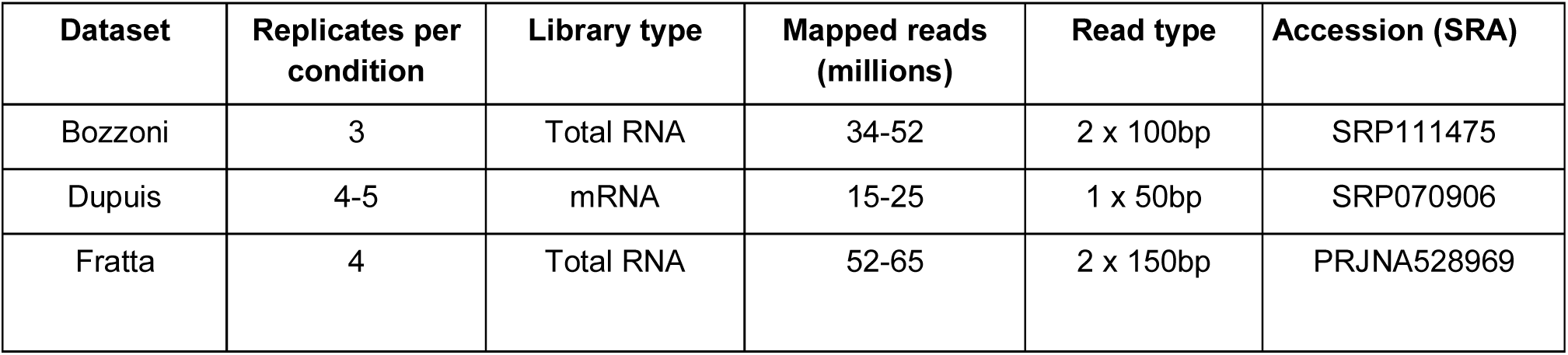
Characteristics of the three sequencing datasets.

| Dataset | Replicates per condition | Library type | Mapped reads (millions) | Read type | Accession (SRA) |
| --- | --- | --- | --- | --- | --- |
| Bozzoni | 3 | Total RNA | 34-52 | 2 x 100bp | SRP111475 |
| Dupuis | 4-5 | mRNA | 15-25 | 1 x 50bp | SRP070906 |
| Fratta | 4 | Total RNA | 52-65 | 2 x 150bp | PRJNA528969 |

**Supplementary Table 2:** The results of the individual and joint gene expression analyses. Numbers refer to the number of genes found to be differentially expressed at FDR < 0.05. Strict overlap between joint models refers to genes with FDR < 0.05 in both models. Relaxed overlap refers to genes with FDR < 0.05 in one model and P < 0.05 in the other.

|  | **Bozzoni**<br>**MUT** | **Dupuis**<br>**MUT** | **Fratta**<br>**MUT** | **Bozzoni**<br>**KO** | **Dupuis**<br>**KO** | **Fratta**<br>**KO** |
| --- | --- | --- | --- | --- | --- | --- |
| Individual analysis | 19 | 1552 | 88 | 100 | 2916 | 151 |
| Joint analysis | **754** |  |  | **2136** |  |  |
| Overlapping joint model | 5 | 368 | 57 | 51 | 1007 | 114 |
| Unique to dataset | 14 | 1184 | 31 | 49 | 1909 | 37 |
| Overlap (strict) | 329 |  | **425** |  | 1711 |  |
| Overlap (relaxed) | 186 |  | **1318** |  | 961 |  |

**Supplementary Table 3:** The results of the individual and joint splicing analyses. Numbers refer to number of splicing events found to be differentially used at FDR < 0.05

|  | **Bozzoni**<br>**MUT** | **Dupuis**<br>**MUT** | **Fratta**<br>**MUT** | **Bozzoni**<br>**KO** | **Dupuis**<br>**KO** | **Fratta**<br>**KO** |
| --- | --- | --- | --- | --- | --- | --- |
| Total (FDR < 0.05) | 31 | 1 | 56 | 211 | 46 | 230 |
| Joint model | **93** |  |  | **890** |  |  |
| Overlapping joint model | 7 | 1 | 30 | 143 | 38 | 169 |
| Unique to dataset | 21 | 0 | 11 | 67 | 8 | 58 |
| Overlap (strict) | 33 |  | **60** |  | 830 |  |
| Overlap (relaxed) | 16 |  | **405** |  | 501 |  |

**Supplementary Table 4:** Genes overlapping with Luisier et al:

| gene<br>name | coords<br>(mm10) | event<br>length/<br>bp | variant<br>type | nearest FUS<br>iCLIP cluster/kb | median<br>phyloP |
| --- | --- | --- | --- | --- | --- |
| *Atp13a3* | chr16:30357274-30361420 | 4146 | complex | 0.54 | 0.234 |
| *Ccdc88a* | chr11:29494093-29499335 | 5242 | complex | 66 | 0.439 |
| *Cdc16* | chr8:13767587-13768561 | 974 | retained intron | 158 | 0.244 |
| *Fbxl5* | chr5:43759825-43760707 | 882 | retained intron | 646 | 0.002 |
| *Fus* | chr7:127972770-127974400 | 1630 | retained intron | 0 | 1.206 |
| *Hnrnpdl* | chr5:100036195-100036481 | 286 | retained intron | 0.31 | 0.061 |
| *Mfn1* | chr3:32562893-32563012 | 119 | retained intron | 0 | 0.255 |
| *Ncor1* | chr11:62401267-62403799 | 2532 | complex | 2.80 | 0.878 |
| *Papola* | chr12:105829277-105834710 | 5433 | complex | 7.32 | 0.234 |
| *Rbm6* | chr9:107833507-107838835 | 5328 | retained intron | 13.8 | 0.150 |
| *Srsf5* | chr12:80947865-80948129 | 264 | retained intron | 0 | 1.712 |
| *Tcerg1* | chr18:42550108-42551110 | 1002 | retained intron | 25.5 | 0.428 |

**Supplementary Table 5:**
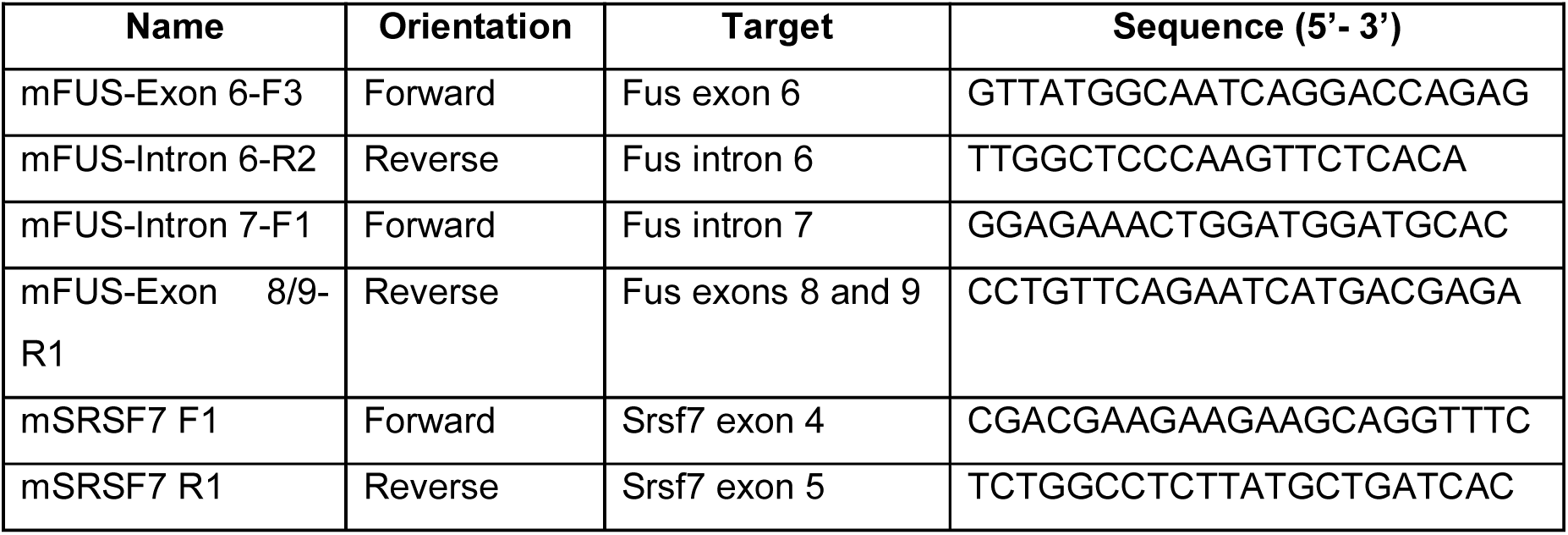
Primer sequences used for RT-PCR in mouse.

**Supplementary Table 6:**
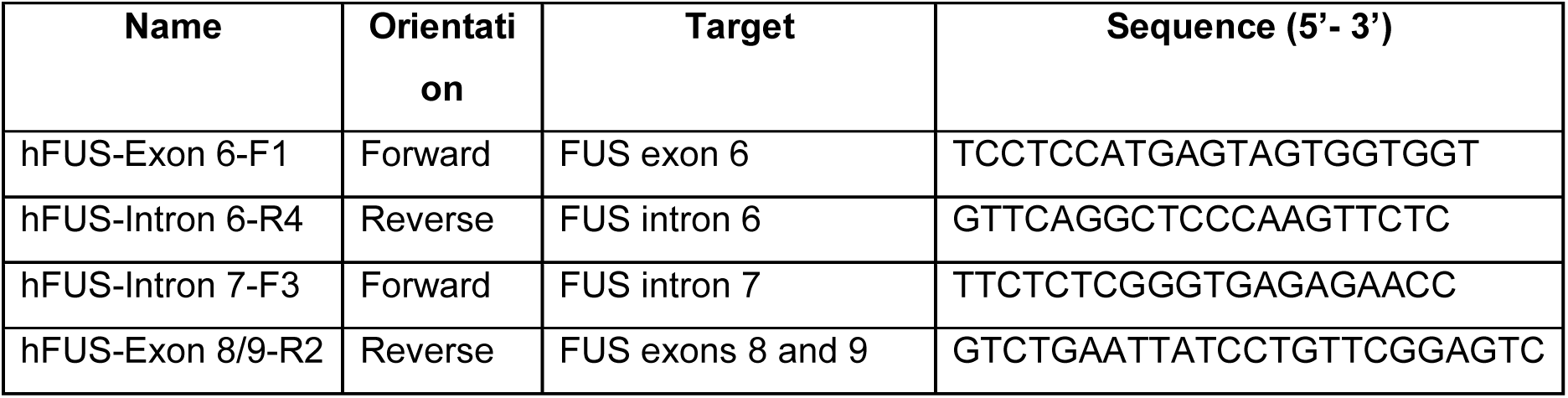
Primer sequences used for RT-PCR in human.

**Supplementary Table 7:**
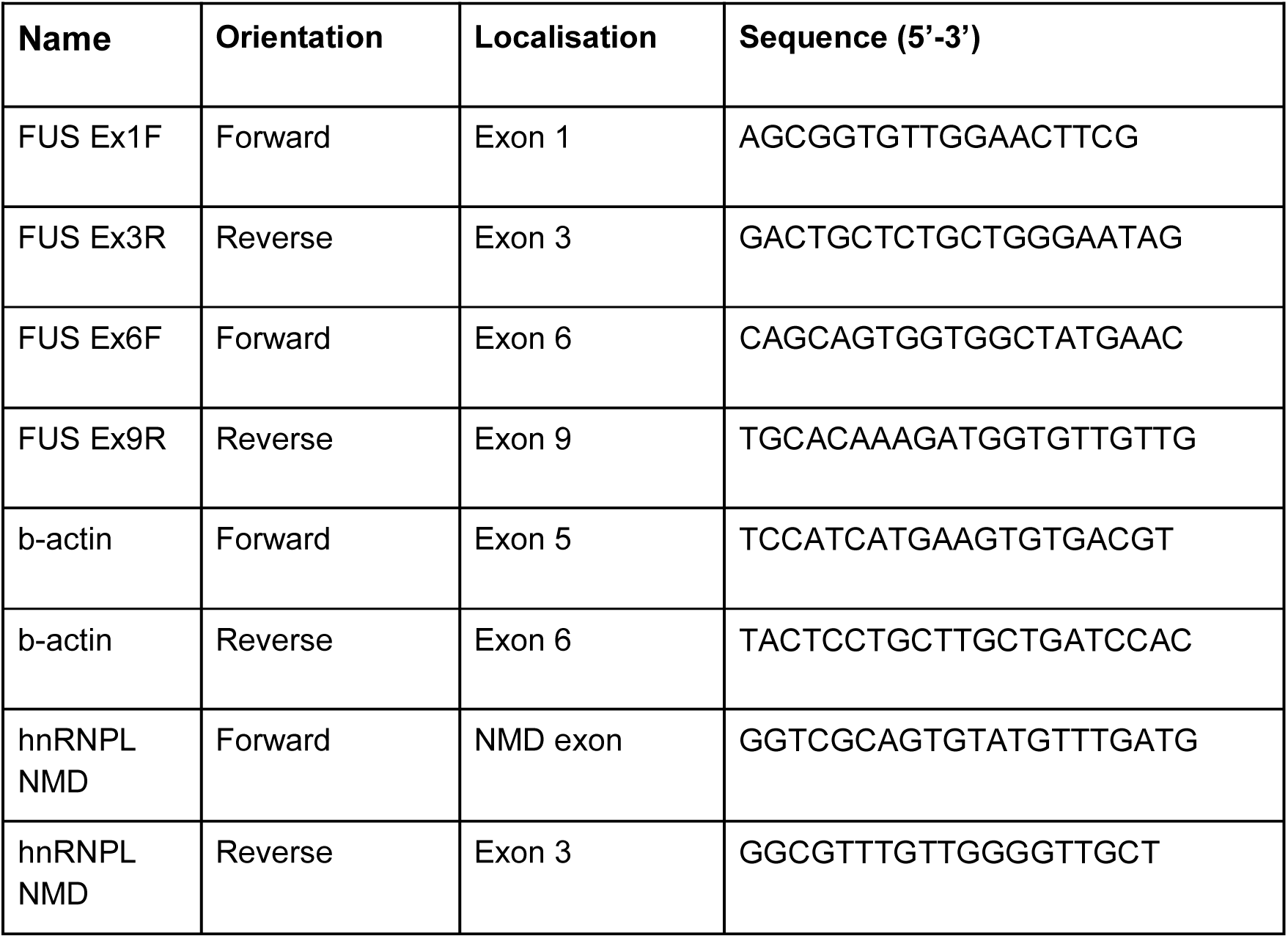
primers used for RT-qPCR in human cells.

**Supplementary Figure 1:**
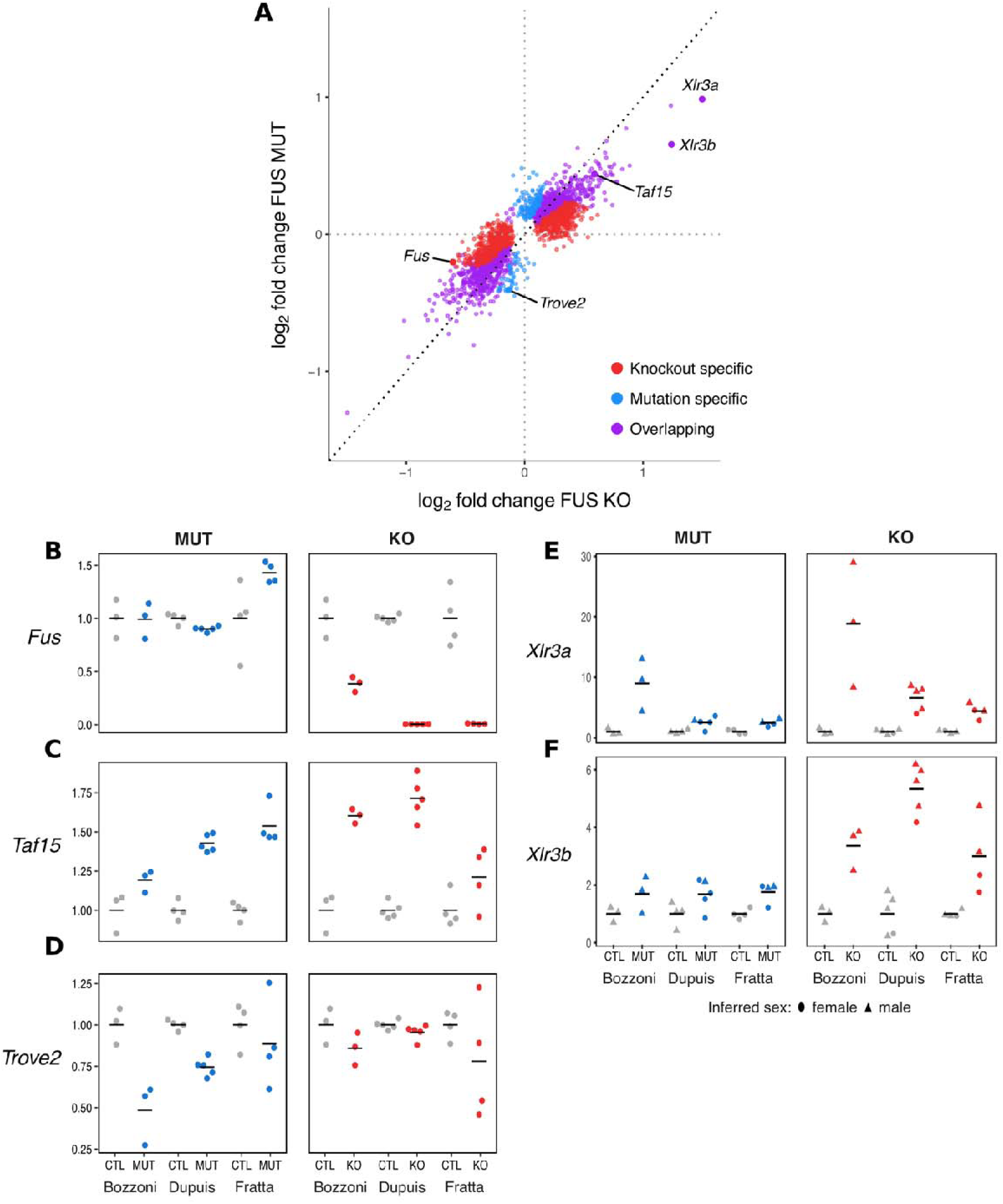
Overlapping gene expression models show concordant effects of KO and MUT. (**A**) Log_2_ fold change of FUS KO plotted against log_2_ fold change in FUS MUT for all three categories of genes. (**B,C,D**) Expression plots for *Fus, Taf15,* and *Trove2* in each sample of each dataset. Library size-normalised expression in MUT and KO samples is normalised to that of each set of wildtype littermates (CTL). (**E,F**) Expression for *Xlr3a* and *Xlr3b.* Inferred sex of the samples from Y chromosome gene expression: females (circles) and males (triangles). Library size-normalised expression in MUT and KO samples is normalised to that of each set of wildtype littermates (CTL).

**Supplementary Figure 2:**
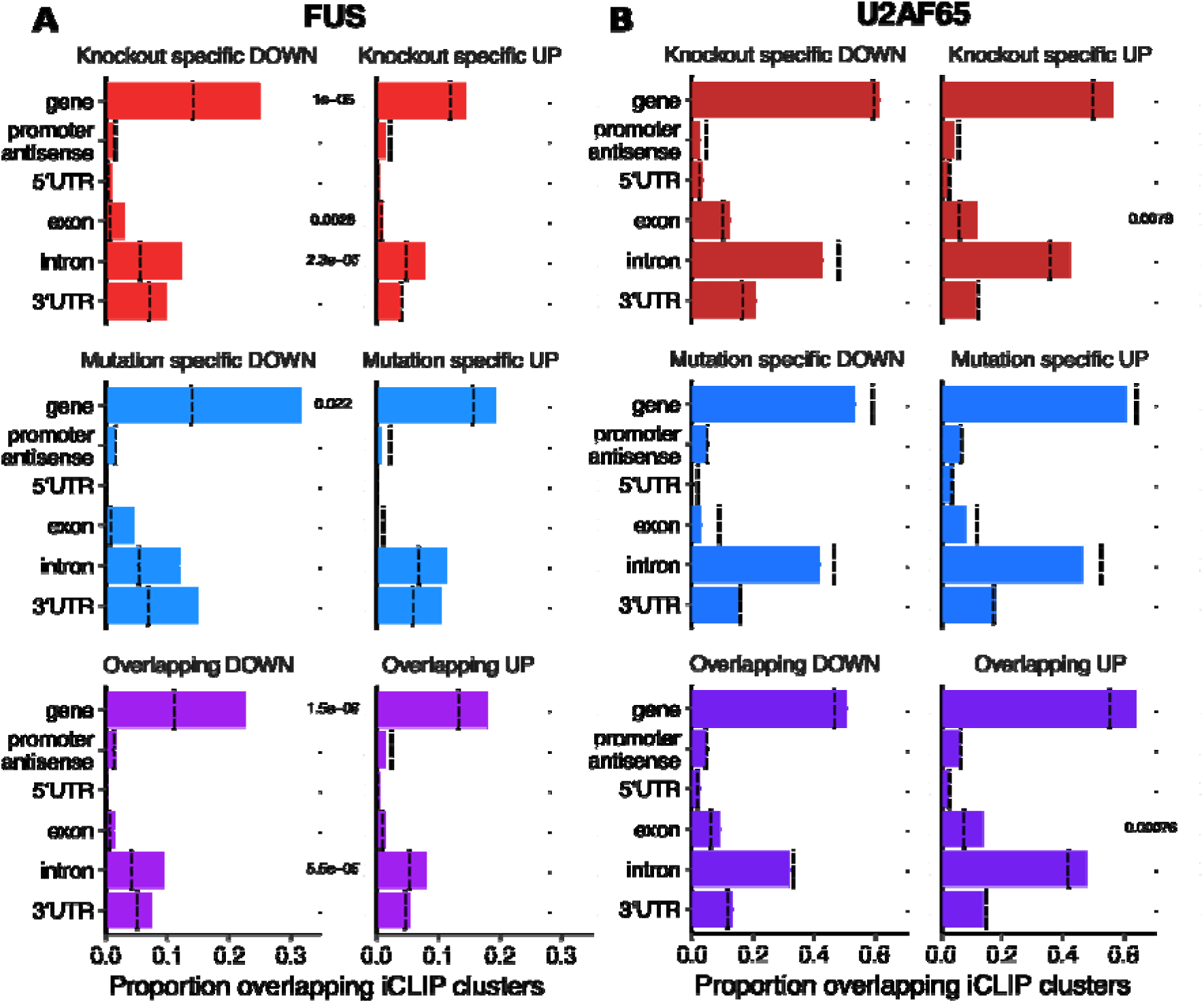
Downregulated genes are enriched in FUS iCLIP clusters. (**A**) Proportion of genes that overlap with a FUS iCLIP cluster. Genes divided by direction of change and group (Knockout-specific, Mutation-specific and Overlapping). Proportions in null sets depicted with black dotted lines. (**B**) As above, but for U2AF65 iCLIP. All P-values corrected for multiple testing with Bonferroni method. Any P-value > 0.05 not shown.

**Supplementary Figure 3:**
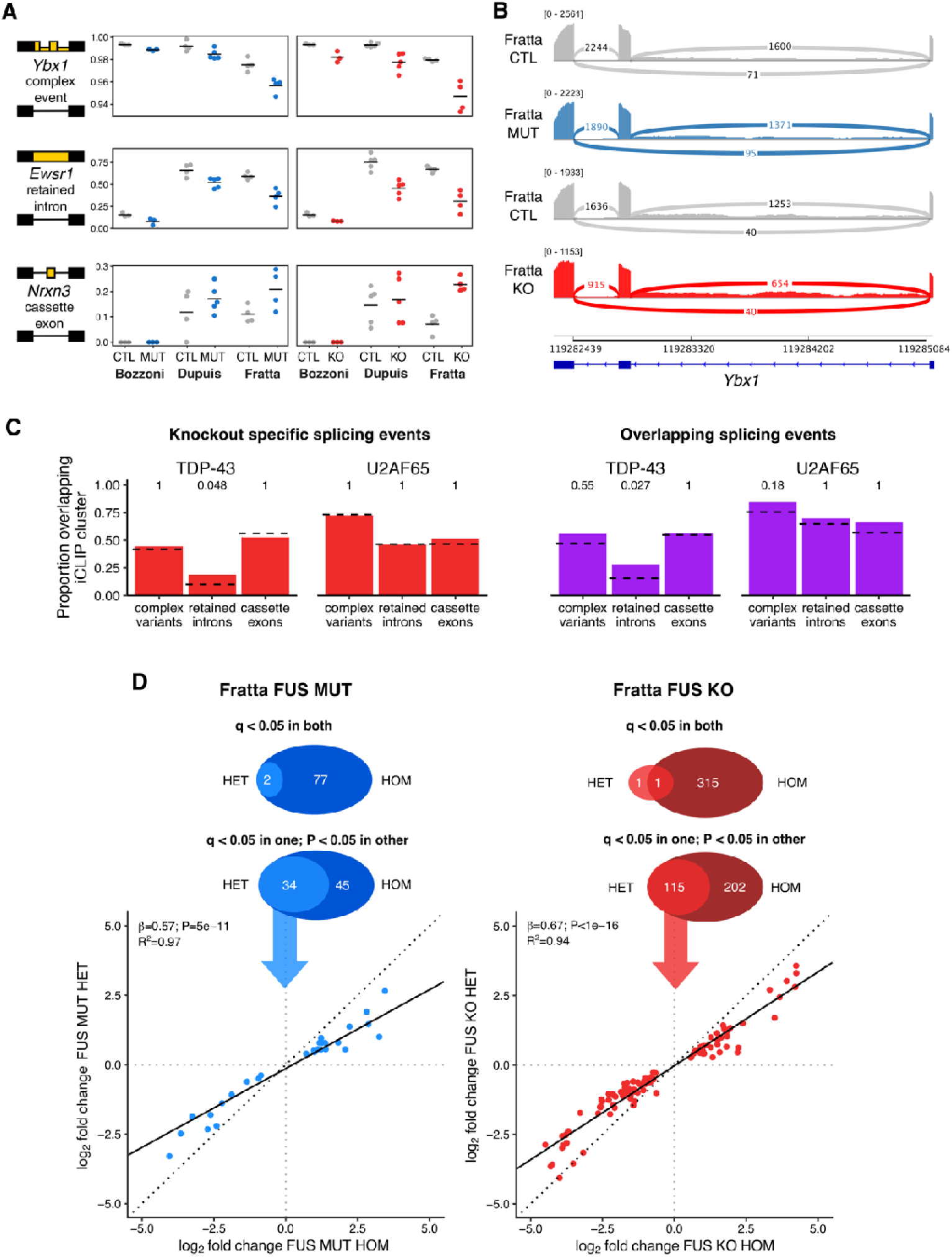
accompaniment to splicing analyses. (**A**) Percentage spliced in (PSI) for three splicing events in the six comparisons. (**B**) Representative RNA-seq traces for a complex event in *Ybx1* in the Fratta MUT and KO samples with their respective controls. (**C**) Proportions of knockout-specific and overlapping splicing events in each category that contain TDP-43 or U2AF65 iCLIP peaks. P-values corrected for multiple testing with Bonferroni method. (**D**) Comparing splicing events found in Fratta MUT and KO homozygous samples with those found in heterozygous samples.

**Supplementary Figure 4:**
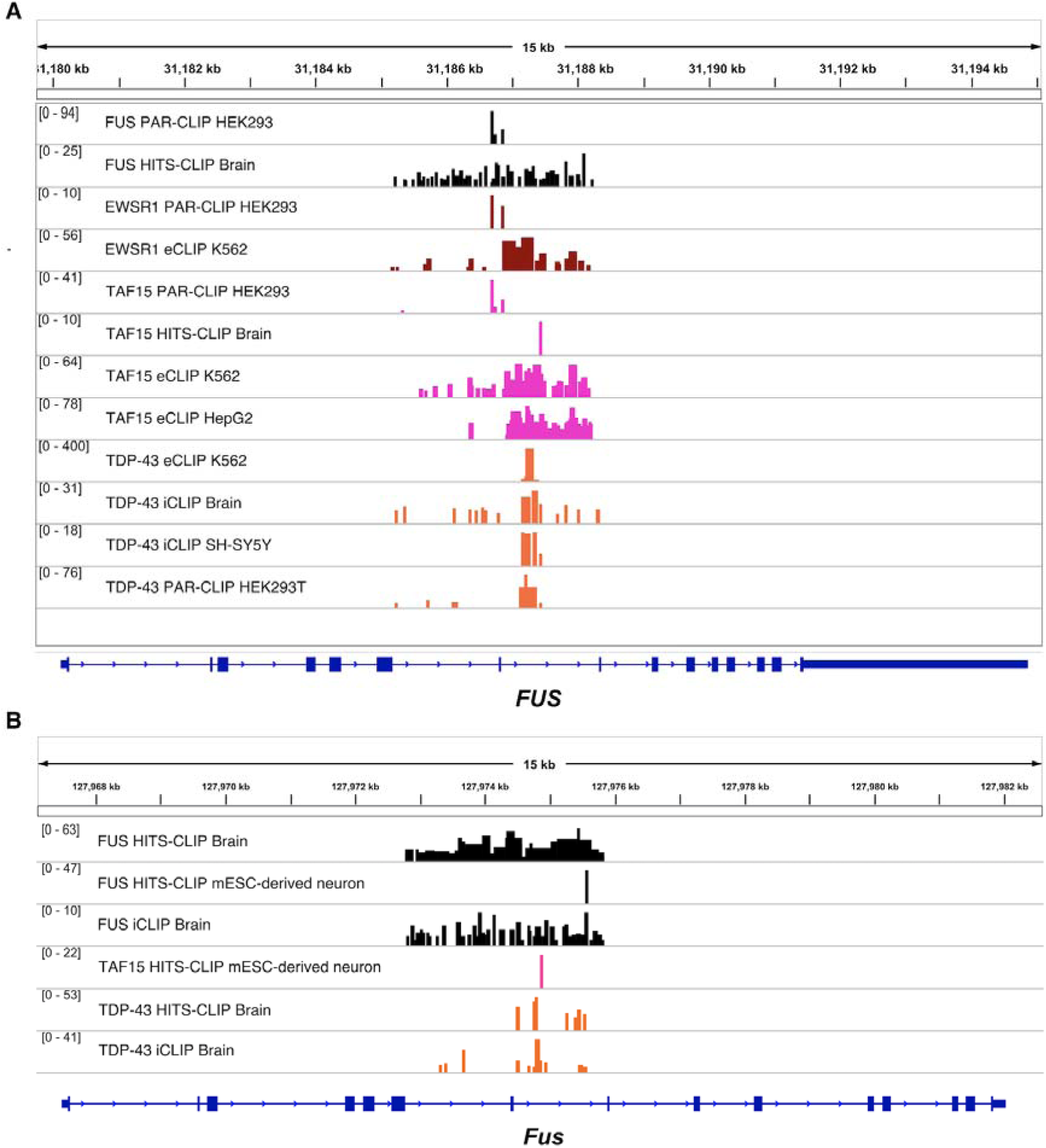
FUS, EWSR1, TAF15 and TDP-43 bind FUS introns 6 and 7 in human and mouse. (**A**) IGV traces across the human *FUS* gene of all CLIP data from the POSTAR/CLIPdb database of the 4 RBPs. Scale indicates the maximum number of reads. (**B**) IGV traces across the mouse *Fus* gene. No mouse EWSR1 CLIP was present in the POSTAR/CLIPdb database.

**Supplementary Figure 5:**
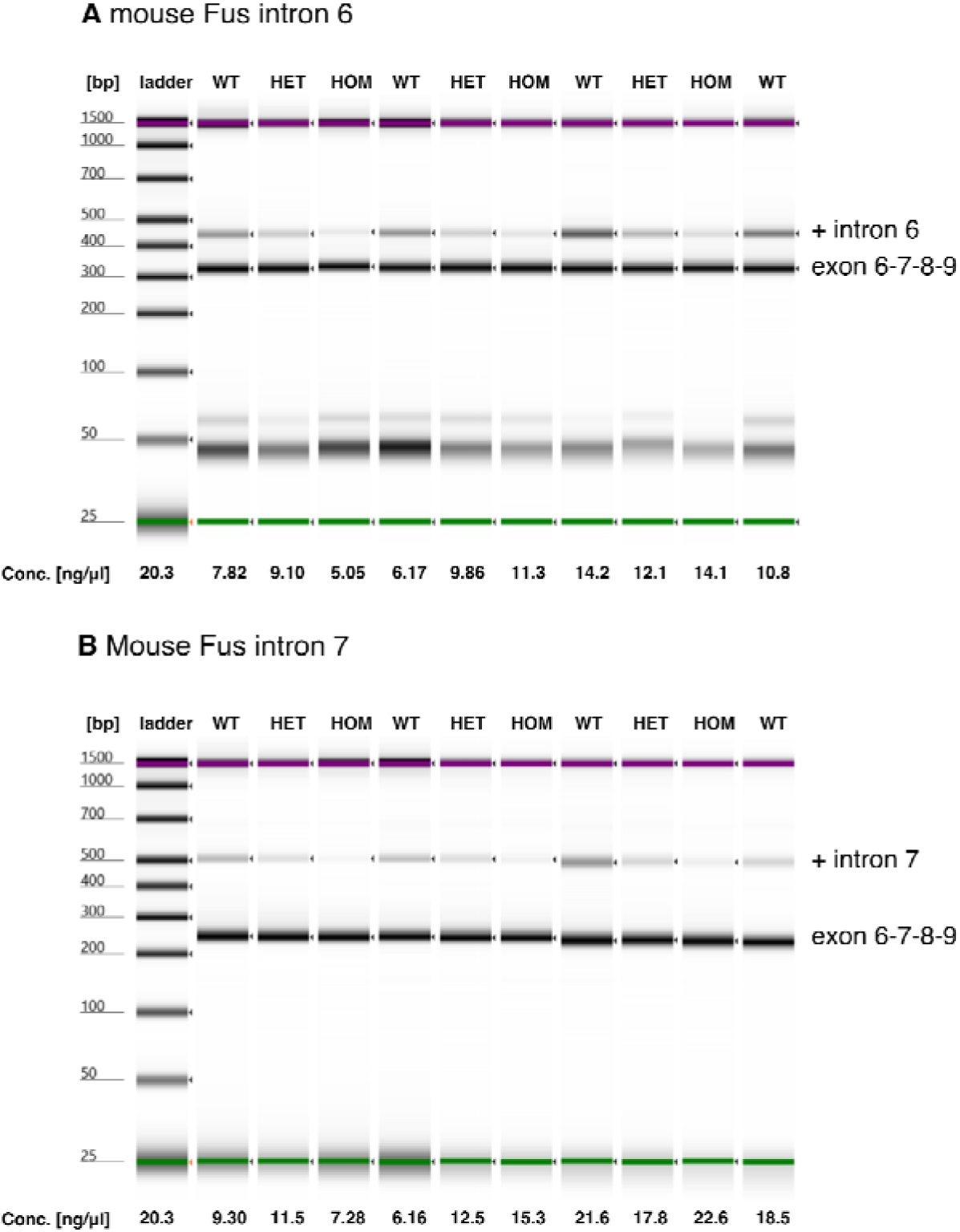
RT-PCR of Fus intron retention in mouse. **(A)** TapeStation traces from RT-PCR with two primers targeting FUS mRNA between exons 6 and 9 and a third primer targeting FUS intron 6. Samples with wildtype Fus (CTL), heterozygous (HET) or homozygous FUS delta14 (HOM). RNA concentrations are taken from the TapeStation itself. Bands used in quantification are annotated. **(B)** As before for Fus intron 7.

**Supplementary Figure 6:**
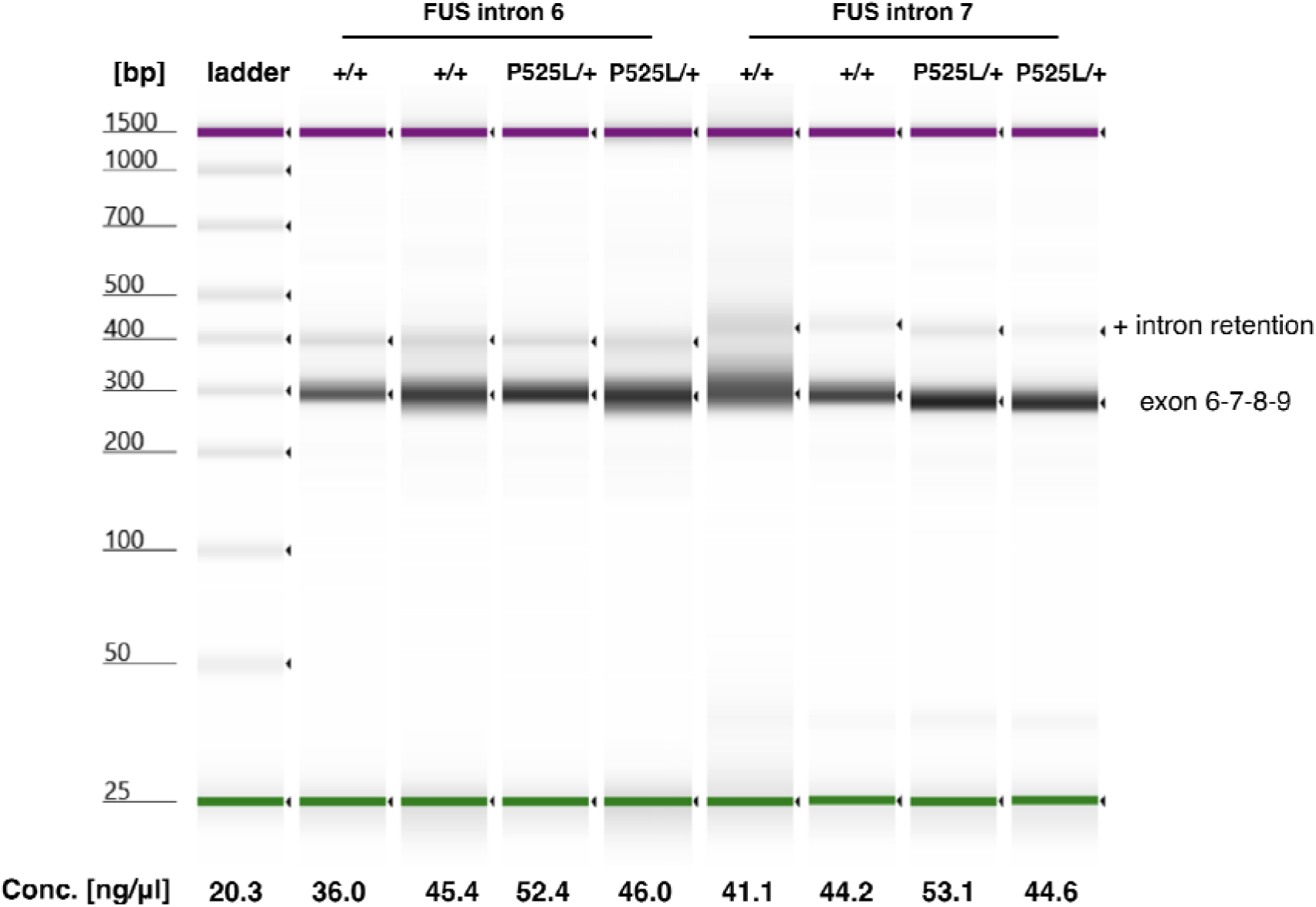
RT-PCR of FUS intron retention in human samples. **(A)** TapeStation traces from RT-PCR with two primers targeting FUS mRNA between exons 6 and 9 and a third primer targeting FUS intron 6 (left 4 lanes) or intron 7 (right 4 lanes). Samples taken from a patient with wildtype FUS (+/+) and a patient heterozygous for the FUS P525L mutation (P525L/+). Replicates are technical, derived from separate RNA extractions. RNA concentrations are taken from the TapeStation itself. Bands used in quantification are annotated. **(B)** As before for Fus intron 7.

**Supplementary Figure 7:**
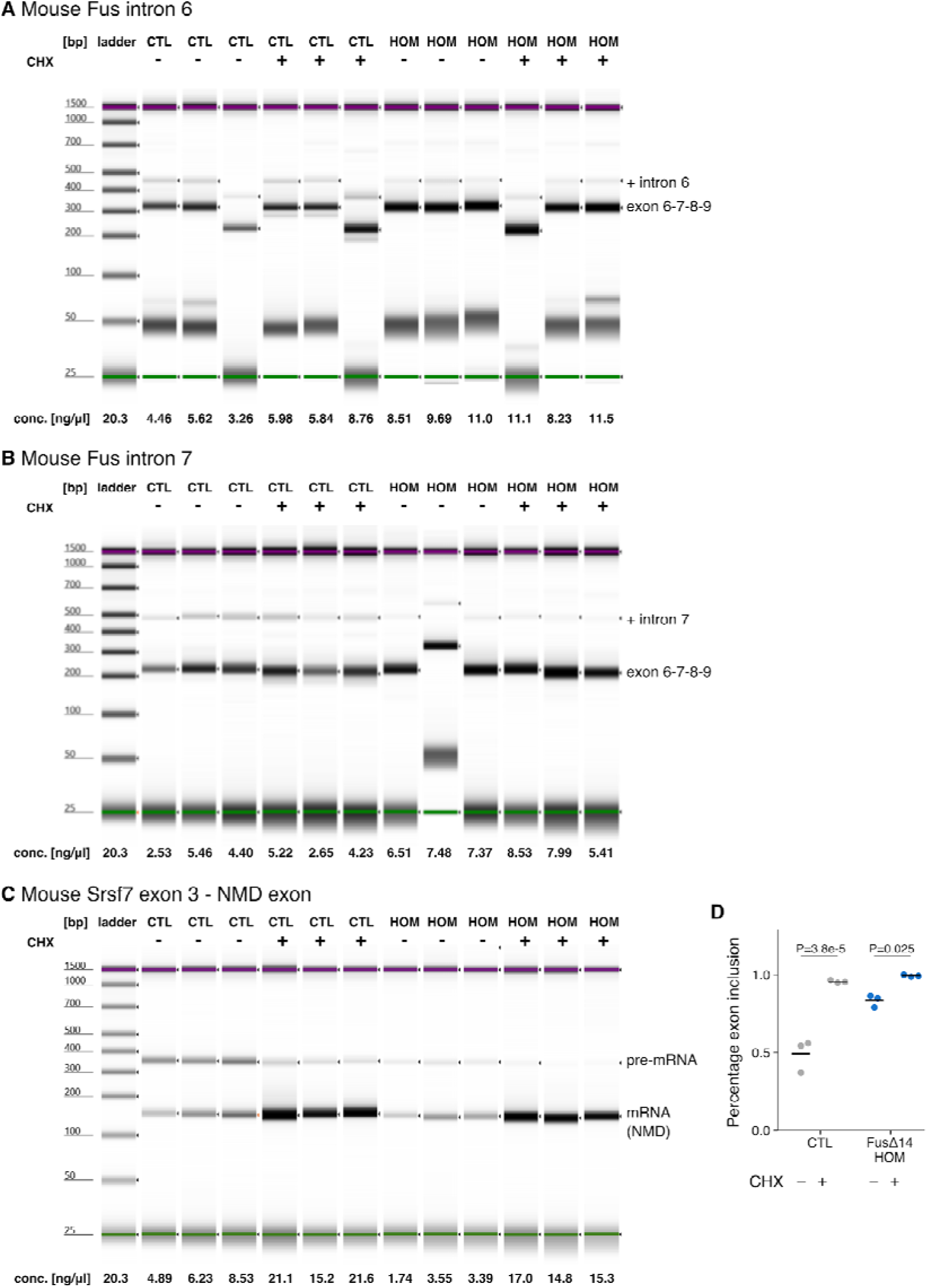
Cycloheximide inhibition experiments. (A) TapeStation traces from RT-PCR with primers targeting Fus intron 6. Samples with wildtype Fus (CTL) or homozygous FUS delta14 (HOM) with or without treatment with Cycloheximide (CHX). RNA concentrations are taken from the TapeStation itself. Bands used in quantification are annotated. (B) As before, but for Fus intron 6. (C) As before but amplifying spliced and unspliced RNA between Srsf7 exon 3 and its known NMD exon. (D) Quantification of Srsf7 mRNA against total. ANOVA treatment P = 1.1e-5; genotype P = 3.7e-4; interaction P = 1.4e-3

**Supplementary Figure 8:**
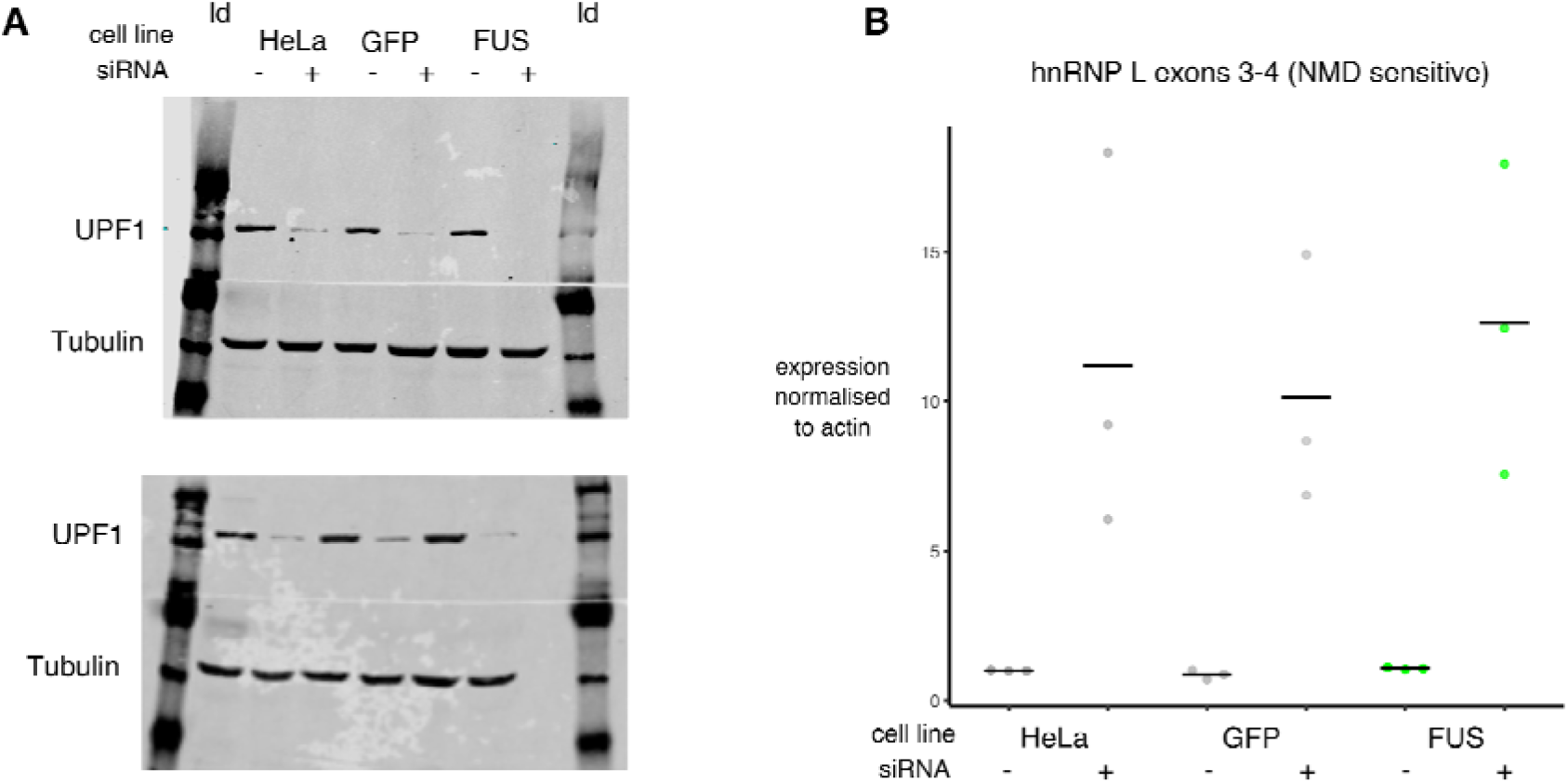
FUS overexpression qPCR and UPF1 knockdown. (**A**) Two independent western blotting experiments demonstrate efficient knockdown of UPF1 protein by siRNA. Tubulin used as a loading control. ld: protein ladder. (**B**) Quantification of expression of hnRNP L NMD-sensitive transcript by RT-qPCR demonstrates UPF1 knockdown is sufficient to inhibit NMD. HeLa - control HeLa cell line; GFP - HeLa cells expressing GFP construct; FUS - HeLa cells expressing codon-optimised FUS transcript. + denotes UPF1 siRNA, - denotes a scrambled siRNA.

**Supplementary Figure 9:**
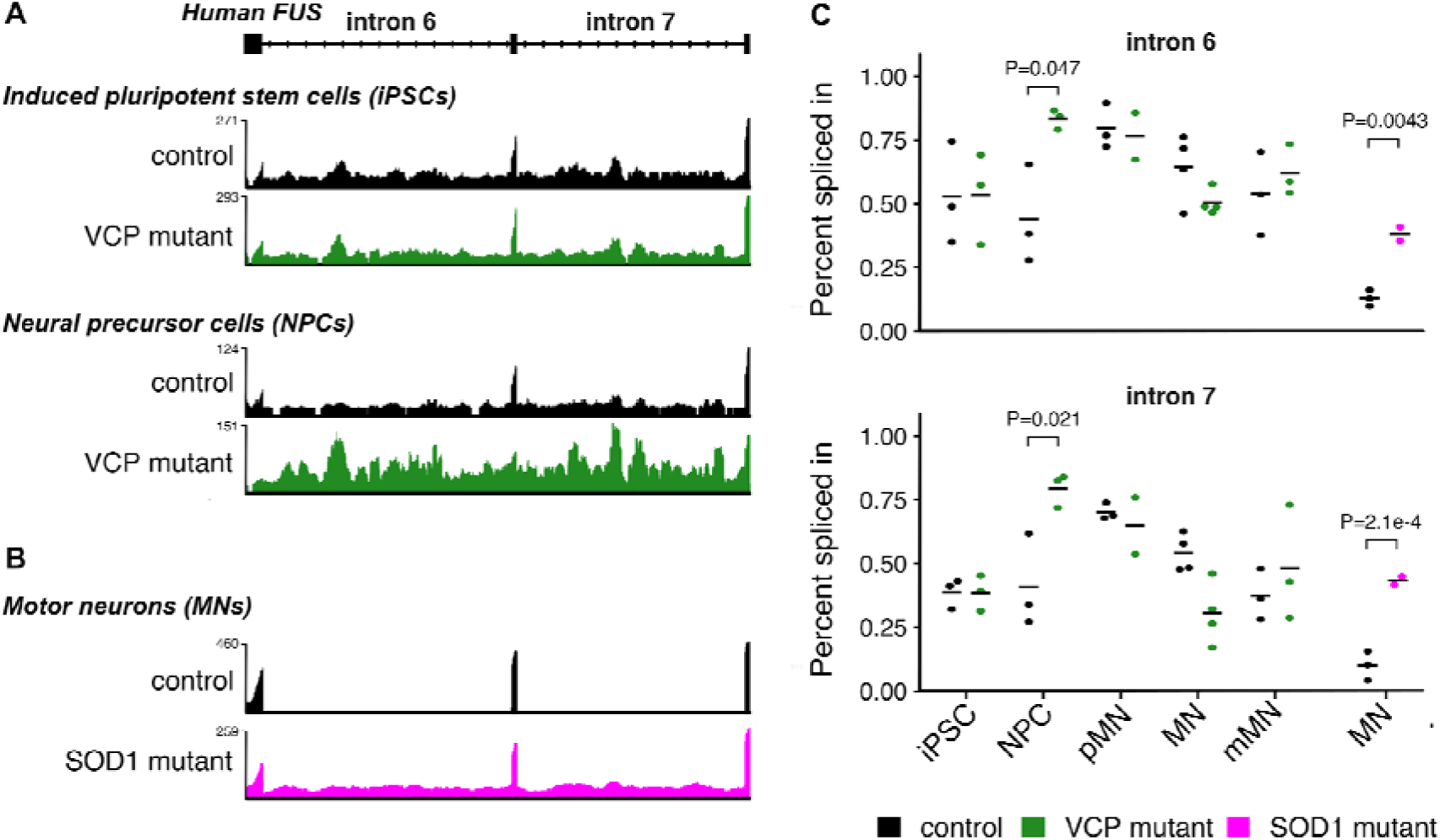
FUS intron retention is dysregulated in different ALS models (A) RNA-seq traces from human induced pluripotent stem cell and neural precursor cells with and without mutations in VCP show a selective increase in retention of introns 6 and 7. (B) RNA-seq traces from motor neurons (MN) with and without the A4V mutation in SOD1. (C) Percent spliced in quantification of all samples across neural differentiation from induced pluripotent stem cell, neural precursor cell (NPC), precursor motor neuron (pMN), immature motor neuron (MN) and mature motor neuron (mMN). ANOVA on PSI ∼ cell type: genotype intron 6 P=0.019 intron 7 P=0.00052. Individual P values from Tukey post-hoc test. For the MN samples with and without SOD1 mutations, P-values were taken directly from the splicing analysis.

## Methods

### Data availability

RNA-seq data for the FUS-Δ14 mutant and FUS KO homozygous and heterozygous spinal cord samples with their respective wildtype littermate controls has been deposited at the Sequence Read Archive (SRA) with project accession number PRJNA528969. FUS-ΔNLS and FUS KO mouse brain samples with their respective controls (Scekic-Zahirovic et al. 2016) were downloaded from SRA accession SRP070906. FUS P517L mutants, FUS KO and shared control samples from mouse motor neurons (Capauto et al. 2018) were downloaded from SRA accession SRP111475. Data from a series of motor neuron differentiation experiments, where induced pluripotent stem cells with and without VCP mutations were differentiated to mature motor neurons with RNA-seq libraries created from cells taken at 0,7,14,21 and 35 days following differentiation (Luisier et al. 2018), were downloaded from the Gene Expression Omnibus with accession GSE98290. Stem cell-derived human motor neurons carrying SOD1 A4V mutations and their isogenic controls (Kiskinis et al. 2014) were downloaded from GSE54409. FUS, U2AF65 and TDP-43 mouse iCLIP (Rogelj et al. 2012) clusters were downloaded from http://icount.biolab.si/. TDP-43, FUS, EWSR1 and TAF15 CLIP data from human and mouse were downloaded from POSTAR (Hu et al. 2017). When the same CLIP sample had been processed with more than one peak caller, the Piranha caller was selected for presentation.

### Mouse lines

FUS Δ14 mice were previously described (Devoy et al. 2017). FUS knockout mice were obtained from the Mouse Knockout Project (Fustm1(KOMP)Vlcg). All procedures for the care and treatment of animals were in accordance with the Animals (Scientific Procedures) Act 1986 Amendment Regulations 2012.

### RNA sequencing

For RNA sequencing experiments FUS Δ14 or KO heterozygous and homozygous mice were compared to their respective wild type littermates. Spinal cords were collected from E17.5 mouse embryos. Tissues were snap frozen, genotyped and total RNA was extracted from the appropriate samples using Qiazol followed by the mini RNAeasy kit (Qiagen). RNA samples used for sequencing all had RIN values of 9.9 or 10. cDNA libraries were made at the Oxford Genomics facility using a TruSeq stranded total RNA RiboZero protocol (Illumina). Libraries were sequenced on an Illumina HiSeq to generate paired end 150bp reads.

### Bioinformatic analysis

Mouse data was aligned to the mm10 build of the mouse genome and human data aligned to the hg38 build of the human genome using STAR (v2.4.2) (Dobin et al. 2013). Prior to differential expression analysis, reads were counted across genes with HTSeq (Anders, Pyl, and Huber 2015). All code for exploratory data analysis, statistical testing and visualisation was carried out in the R statistical programming language (Gentleman and Ihaka 1996), using the tidyverse suite of packages (Wickham 2017). Data visualisation and figure creation was aided by the patchwork (https://github.com/thomasp85/patchwork), stargazer (Hlavac 2015), ggbeeswarm (Clarke and Sherrill-Mix 2017) and ggrepel packages (Slowikowski, n.d.). All R code written for the project is publicly available as interactive Rmarkdown notebooks (https://github.com/jackhump/FUS_intron_retention).

### Joint modelling of differential expression

Each dataset consists of FUS knockout samples, FUS NLS mutation samples and wildtype controls. In the Bozzoni dataset the controls are shared but in the other two datasets the knockout and mutation samples have their own separate controls for use in two-way comparisons. Differential gene expression was tested with DESeq2 (Love, Huber, and Anders 2014). Initially each comparison (wildtype vs knockout or wildtype vs mutation) was performed separately for each dataset, creating six individual analyses. To boost power and create a set of high confidence changes, two joint models were created using either the knockout (KO) or mutation (MUT) samples with their specific controls. The joint model uses all the samples of the same comparison together in a general linear model with a dataset-specific covariate. DESeq2 uses a Bayesian shrinkage strategy when estimating the log2 fold change. For each gene the estimated log2 fold change is a combination of the three individual datasets. Genes are reported as significantly differentially expressed at a false discovery rate (FDR) threshold of 0.05 (Benjamini and Hochberg, 1995). For plots, gene expression values are raw counts multiplied by each sample’s size factor generated by DESeq2. These normalised counts are then normalised to the wildtype samples for each dataset to visualise the relative change in expression.

To assess the level of overlap between the KO and MUT joint models, two different overlap thresholds were employed. The first, a more conservative threshold, depends on a gene being significant at FDR < 0.05 in both datasets. The second, more relaxed threshold, calls a gene as significant if it falls below FDR < 0.05 in one dataset and has an uncorrected P-value < 0.05 in the other.

### Joint modelling of splicing

SGSeq was run on all the samples together to discover and classify all potential splicing events using the default parameters for finding both annotated and novel splicing (Goldstein et al., 2016). Differential splicing for individual comparisons and joint models with a dataset-specific covariate were performed using DEXSeq (Anders et al., 2012). The same overlap threshold strategies were employed as for differential gene expression. SGSeq looks for all potential splicing events in each sample and then counts the reads supporting either the inclusion or exclusion of that splicing variant. Percentage Spliced In (PSI) values (Katz et al., 2010) for each splicing variant in each sample were calculated by taking the read counts supporting the inclusion event and dividing by the total reads in that event.

### Functional analysis of genes and splicing events

iCLIP data on FUS and U2AF65 from mouse brain (Rogelj et al., 2012) was reprocessed by the iCOUNT iCLIP analysis pipeline (http://icount.biolab.si/), and the set of FUS iCLIP clusters that passed enrichment against background at FDR < 0.05 were downloaded. Only iCLIP clusters with a minimum of two supporting reads were kept. Untranslated region (UTR) and coding exon (CDS) annotation were taken from GENCODE mouse (comprehensive; mouse v12). Any intron retention, nonsense mediated decay or “cds end nf” transcripts were removed. UTR coordinates were split into 5’ and 3’ UTR based on whether they overlapped an annotated polyadenylation site or signal (GENCODE mouse v18 polyadenylation annotation). 3’ UTRs were extended by 5 kilobases downstream to capture any unannotated sequence. Introns were defined as any gaps in the transcript model between CDS and UTR-antisense coordinates were taken by flanking the 5ÚTR sequence by 5kb upstream and inverting the strand. Overlaps between iCLIP clusters and genomic features were created for each set of differentially expressed genes, split into upregulated (log_2_ fold change > 0) or downregulated (log_2_ fold change < 0). Overlaps were done in a strand-specific manner, with only iCLIP clusters in the same direction being used.

Whether an iCLIP cluster overlaps a genomic region depends on both the affinity of the chosen protein for RNA sequence of the motif and the abundance of the RNA in the cell (Chakrabarti et al. 2018). In addition, a longer region would be more likely to overlap an iCLIP cluster by random chance than a shorter region. When comparing sets of genomic regions, whether genes or splicing events, this must be taken into account. See appendices for distributions of lengths and expressions between significant and non-significant genes and splicing events.

To test for enrichment of FUS iCLIP clusters in upregulated and downregulated genes, each set of tested genes was compared to a set of null genes with no evidence of differential expression (P > 0.05 in both models). The null set was then restricted to genes with both length and expression values that were within the first and third quartile of those of the test gene set. The expression values were calculated by taking the mean number of reads covering each gene in the Fratta wildtype samples, with each sample read count first normalised by the library size factor for each sample calculated by DESeq2. The proportion of each set of genes overlapping an iCLIP peak was then compared to that of the null set with a χ^2^ test of equal proportions.

For the splicing events found in the joint models, enrichment tests were performed for different genomic features. For these tests the coordinates of the entire encompassing intron were used for each splicing variant. Each test set of splicing events was compared to a matched set of null splicing events where P > 0.05 in both joint models. The null events were chosen to have length and expression levels within the first and third quartiles of that of the test set. Proportions of overlap with ICLIP clusters between splicing events and the null set were tested using a χ^2^ test of equal proportions. As a positive control in both analyses, the same overlaps were computed with iCLIP clusters from U2AF65, also from (Rogelj et al. 2012).

Per nucleotide phyloP conservation scores (Pollard et al. 2010) comparing mouse (mm10) with 60 other vertebrates was downloaded from UCSC. The median phyloP score was calculated for each splicing variant and compared.

### RT-PCR - intron retention validation

Primers were designed using Primer3 (Koressaar and Remm 2007) and in silico PCR (UCSC). For both human and mouse FUS, the forward primer was designed for exon 6 and the reverse primer designed to span the spliced exon 8/9 junction to preferentially amplify spliced FUS mRNA. An additional third primer was designed to amplify a section of either intron 6 or intron 7. Primer sequences are listed in Supplementary Table 5.

Cells were obtained from mouse spinal cord and/or cultured mouse embryonic fibroblasts resuspended in Trizol (Thermo Fisher). RNA was extracted using miRNeasy Mini Kit (Qiagen) following the manufacturer’s instructions. cDNA was obtained from extracted RNA using SuperScript IV Reverse Transcriptase kit (Thermo Fisher). Briefly, a mix was made of RNA template (500ng for mouse brain; 100ng for cultured cells (cycloheximide treatment)), 10mM dNTP, 50mM oligo d(T)20, 50mM random hexamer followed by 5 min of incubation at 65°C and 1 min in ice. Mix was then complemented with 5X SuperScript IV Reverse Transcriptase buffer, 100nM DTT, RNase OUT and SuperScript IV Reverse Transcriptase buffer followed by incubation at 23°C, 55°C and 80°C, 10 min each.

RT-PCR was carried out using 10X AccuPrime Taq DNA polymerase mastermix system (Invitrogen). Each PCR reaction mix contained 5ng of gDNA, 10mM of forward and reverse primers. cDNA was amplified with the following conditions: Intron 6 retention: One cycle of 5 min at 95°C, followed by 30 cycles of 30 sec at 95°C, 30 sec at 56°C, and 30 sec at 68°C, and finishing with 5 min incubation at 68°C. Intron 7 retention: One cycle of 5 min at 95°C, followed by 30 cycles of 30 sec at 95°C, 30 sec at 61°C, and 30 sec at 68°C, and finishing with 5 min incubation at 68°C. Srsf7 NMD positive control: One cycle of 5 min at 95°C, followed by 35 cycles of 30 sec at 95°C, 30 sec at 58°C, and 15 sec at 68°C, and finishing with 5 min incubation at 68°C. Amplified products were finally obtained using Agilent 4200 TapeStation System following the manufacturer’s instructions. Results were analysed on TapeStation analysis software (Agilent). Intron retention events are plotted as the percentage of integrated area of band corresponding to intron retention. One- or two-way ANOVA designs were employed with pairwise t-tests with Holm correction for multiple testing. For the RT-PCR on human fibroblasts, four technical replicates were obtained from two independent cell cultures, performed at different time point and derived from the same original two human samples (referred as +/+ and P525L/+).

### Cycloheximide treatment and fractionation

Mouse embryonic fibroblasts were treated with 100ug/ml cycloheximide (Sigma) for 6 hours before RNA was extracted with Trizol (Thermo Fisher) and RT-PCR performed as before. As a positive control, primers targeting the NMD-sensitive exon 4 of Srsf7 were used from (Edwards et al., 2016). Nuclear and cytoplasmic fractions were obtained from primary MEF cells using the Norgen Kit following manufacturer’s instructions.

### Plasmids

pLVX-EF1a-TS-EGFP-IRES-Puro was cloned by introducing an N-terminal Twin-Strep (TS)-tagged EGFP cDNA (DNA String byGeneArt, Life Technologies) into the EcoRI and BamHI sites of pLVX-EF1a-IRES-Puro (Clontech, Cat. Nr. 631988). pLVX-EF1a-TS-OPT-FUS-IRES-Puro was cloned by introducing an N-terminal Twin-Strep (TS)-tagged codon optimized FUS cDNA (Gene synthesis by GeneArt, Life Technologies) into the EcoRI and BamHI sites of pLVX-EF1a-IRES-Puro (Clontech, Cat. Nr. 631988).

### Stable cell line generation

293T cells were cultured in DMEM/F12 supplemented with 10% heat-inactivated, tetracycline-free foetal calf serum (FCS) (Contech, Cat. Nr. 631105), penicillin (100 g/ml) (Amimed, Bioconcept Cat. Nr. 4-01F00-H). One day prior to transfection approximately 5×106 HEK293T cells were plated in 150cm2 flasks. 28mg of the pLVX-EF1a vectors and 144ml of the fourth generation Lenti-X HTX Packaging Mix (Clontech, Cat. Nr. 631249) were transfected using the Xfect transfection reagent (Clontech, Cat. Nr. 631317). 24 hours post transfection the medium was exchanged. 48, 72, and 96 hours post transfection viral particle containing supernatants were harvested and filtered through a 0.45μm PES syringe filter (Membrane Solutions, Cat. Nr. SFPES030045S) followed by a 6-fold concentration using Lenti-X-Concentrator (Clontech, Cat. Nr. 631232) according to the manufacturer’s instructions.

One day before transduction, 2×105 HeLa cells were seeded in four wells of a six-well plate. The next day, the cells were exposed to 1ml concentrated viral supernatant in a total volume of 2ml DMEM+/+ supplemented with 10ug/ml Polybrene (Sigma Aldrich, Cat. Nr. 107689) to increase lentiviral transduction efficiency. The following two days, the same procedure was carried out with virus from the 2nd and 3rd harvest respectively. Finally, the transduced cells were expanded under constant puromycin selection at 2μg/ml.

### Knockdown of UPF1 by siRNAs

Knockdown of UPF1 was carried out in three different HeLa cell lines: wildtype cells, cells containing stably integrated GFP reporter gene, and cells containing stably integrated codon-optimised FUS reporter gene. Knockdown was achieved using siRNAs for UPF1 (GAUGCAGUUCCGCUCCAUUdTdT) and scrambled control sequence (AGGUAGUGUAAUCGCCUUGdTdT, Microsynth, CH). In short, 2-3 x 105 cells were seeded into a 3.5-cm dish and transfected the following day with 40 nM siRNAs using Lullaby (OZ Biosciences) according to the manufacturer’s protocol. After 48 hours, cells were re-transfected with 40 nM siRNAs and harvested 48 hours after the second siRNA transfection. Until harvest, cells were split to avoid overgrowth of the cell culture. The efficiency of the knockdown was assessed by western blotting.

### RT-qPCR

RNA analysis was performed according to (Nicholson, Joncourt, and Mühlemann 2012). Briefly, harvested cells were lysed in Trizol reagent (Thermo Fisher) and RNA was isolated according to standard protocol. Prior to reverse transcription (RT), DNase treatment was performed using Turbo DNA-free kit (Invitrogen) to avoid any DNA contamination. cDNA was synthesised using AffinityScript Multiple Temperature Reverse Transcriptase (Agilent) and RT control samples (without addition of RT) were included for each sample. The cDNA was measured in triplicates by RT-qPCR (reaction volume 15 l) using Rotor-Gene Q (Qiagen) and Brilliant III Ultra-Fast SYBR Green qPCR Master Mix (Agilent). Oligonucleotides (final concentration 0.6 M) used in the qPCR measurements are listed in Supplementary Table 8. Ct values were converted to fold changes using the delta-delta-Ct method (Livak and Schmittgen 2001) in R.

### Supplementary Data

1. *joint_model_expression.csv* - Gene expression joint model output
2. *expression_GO_full.tsv* - all GO terms found in each category of differentially expressed genes
3. *joint_model_splicing.csv* - Splicing joint model output with iCLIP distances and phyloP conservation
4. *splicing_GO_full.tsv* - all GO terms found in each category of splicing event

## Acknowledgments

JH was funded by an MRC PhD studentship. PF is funded by an MRC/MNDA LEW Fellowship, the NIHR-UCLH Biomedical Research Centre and the Rosetrees Foundation. GS is funded by a WT Senior Investigator Award (107116/Z/15/Z), Horizon 2020 Research and Innovation programme (739572). GS, MDR, and AI were supported by the UK Dementia Research Institute. This project was also supported by a donation of the Fondation Dufloteau to MDR. We thank Raphaelle Luisier for providing lists of intron retention events.

## Author Contributions

JH and PF designed and conceived the study. NB carried out RNA-sequencing experiments with assistance from CB. JH performed all computational analysis with input from SJ, VP and PF. Validation experiments were performed by CM, DR, MB, MGG, ABE and AD with input from AU. Projects were supervised by AR, IB, MDR, OM, AI, EF, GS, VP, and PF. The paper was written by PF and JH with major contributions from NB and AU, and suggestions from all other authors.

